# Three bacterium-plasmid golden combinations facilitate the spread of ST11/CG258 carbapenemase-producing *Klebsiella pneumoniae* in China

**DOI:** 10.1101/2021.04.21.440730

**Authors:** Cuidan Li, Xiaoyuan Jiang, Tingting Yang, Yingjiao Ju, Zhe Yin, Liya Yue, Guanan Ma, Xuebing Wang, Ying Jing, Xinhua Luo, Shuangshuang Li, Xue Yang, Fei Chen, Dongsheng Zhou

**Affiliations:** CAS Key Laboratory of Genome Sciences & Information, Beijing Institute of Genomics, Chinese Academy of Sciences, China National Center for Bioinformation, Beijing, China; State Key Laboratory of Pathogen and Biosecurity, Beijing Institute of Microbiology and Epidemiology, Beijing, China; University of Chinese Academy of Sciences, Beijing, China; State Key Laboratory of Pathogenesis, Prevention and Treatment of High Incidence Diseases in Central Asia, Xinjiang, China; Beijing Key Laboratory of Genome and Precision Medicine Technologies, Beijing, China

**Keywords:** *Klebsiella pneumoniae*, drug resistance, carbapenemase, plasmid, genomic epidemiology

## Abstract

Carbapenemase-producing *Klebsiella pneumoniae* (cpKP) poses serious threats to public health. Previous studies showed that only ST11/CG258-cpKP successfully disseminated in China, however, the underlying genetic bases are still unknown. We conducted a comprehensive genomic-epidemiology analysis on 420 cpKP isolates from 70 hospitals in 24 Chinese provinces during 2009-2017 based on short-/long-reads sequencing. Three ‘golden’ combinations of host––*bla*_KPC_-carrying plasmids (Clade 3.1+3.2—IncFII_pHN7A8_, Clade 3.1+3.2—IncFII_pHN7A8_:IncR, Clade 3.3—IncFII_pHN7A8_:Inc_pA1763-KPC_) endowed cpKP with advantages both in genotypes (strong-correlation/co-evolution) and phenotypes (resistance/growth/competition), thereby facilitating nationwide spread of ST11/CG258-cpKP. Intriguingly, Bayesian skyline illustrated that the three ‘golden’ combinations might directly lead to the strong population expansion during 2007-2008 and subsequent maintenance of the dissemination of ST11/CG258-cpKP after 2008. We tested drug-resistance profiles and proposed combination treatment regimens for CG258/non-CG258 cpKP. Our findings systematically revealed the molecular-epidemiology and genetic-basis for dissemination of Chinese ST11/CG258 cpKP and reminded us to monitor the ‘golden’ combinations of cpKP- plasmid closely.

## Introduction

Antimicrobial-resistant *<u>K</u>lebsiella <u>p</u>neumoniae* (KP) is listed as the ‘K’ in ESKAPE pathogens, the six most significant and dangerous causes for antimicrobial-resistant nosocomial infections, and has been recognized as a major threat to global public health (1). Carbapenems (e.g. imipenem and meropenem) are the first choice for treatment of severe or refractory infections caused by KP, but clinical carbapenem-resistant KP isolates spread worldwide in recent years (2). It remains the top priority in antimicrobial-resistant *K. pneumoniae* due to high morbidity and mortality, limited treatment options, prolonged hospitalization, and high treatment costs (3–6). The Centers for Disease Control and Prevention (CDC) has classified it as one of the most “urgent” public health threats in the United States (7); according to the report of China Antimicrobial Resistance Surveillance System (CARSS), the resistant rate for the two main carbapenem antibiotics (imipenem and meropenem) has been steadily increasing from 2005 to 2017 (about 20%) (http://www.chinets.com).

Production of exogenous carbapenemases is one of the major causes for carbapenem resistance in KP, and <u>c</u>arbapenemase-producing KP (cpKP) has emerged as a threatening epidemic pathogen in hospital settings (3, 8). The carbapenemase genes in cpKP mainly include *bla*_KPC_, *bla*_NDM_, and *bla*_IMP_, of which .*bla*_KPC_ is the most clinically significant one in most countries (3, 7). They are typically carried on the plasmids of many incompatibility (Inc) groups such as IncFII, X, I, C, N, R, P-2, U, W, and L/M(3, 9). *bla*_KPC_ on these plasmids are usually located on Tn*4401b* and its derivatives in European and American countries (9), but on Tn*6296* and its derivatives in China (10). Tn*4401b* and Tn*6296* are genetically wildly divergent, but both belong to Tn*3*-family unit transposons (9, 10).

Genomic studies showed that the clonal group CG258 is highly associated with cpKP isolates, especially *bla*_KPC_-carrying cpKP (11–15). CG258 mainly consists of ST258 and its single-locus allelic variants ST11 and ST512. ST258 is a recombined hybrid from ST11 and ST442, which contribute to the genome composition of ST258 by 80% and 20%, respectively (16). The cpKP isolates of ST258 and ST512 are mostly prevalent in American and European countries (9, 13, 17), while those of ST11 are highly dominant 3in China (18–20).

On the one hand, there are several publications to explore why ST258 cpKP successfully spread in USA and European countries (8, 21). On the other hand, although 115 STs of cpKP have been identified worldwide so far (9), it is reported that only ST11/CG258 has been successfully clonal spread in China (18–20). What is the cause of this phenomenon? Previous research only reported this phenomenon without deep mining the genetic basis (18–20). A large-scale genomic study on cpKP isolates is necessary to uncover the genetic basis for its dissemination in China, however, it is still lacking until now.

In this study, we employed second- and third-generation sequencing technologies to comprehensively analyze the genomic epidemiology in 420 clinical cpKP isolates collected from multicenter hospitals of 24 provinces of China from 2009 to 2017. The results displayed a panoramic population snapshot of cpKP isolates harboring mainly *bla*_KPC_-carrying plasmids of diverse Inc groups and further provided essential insights into the evolution of host KP–– *bla*_KPC_-carrying plasmids and their role in facilitating the nationwide spread of ST11/CG258 cpKP.

## Results

### Genetic diversity of CG258 and non-CG258 cpKP from China

All the 420 cpKP isolates were sequenced using Illumina platform, and 69 of them were determined for complete genome sequences using PacBio (35/69) or Nanopore (34/69) sequencing platforms (Figure 1 and Table S1).

**Fig. 1.**
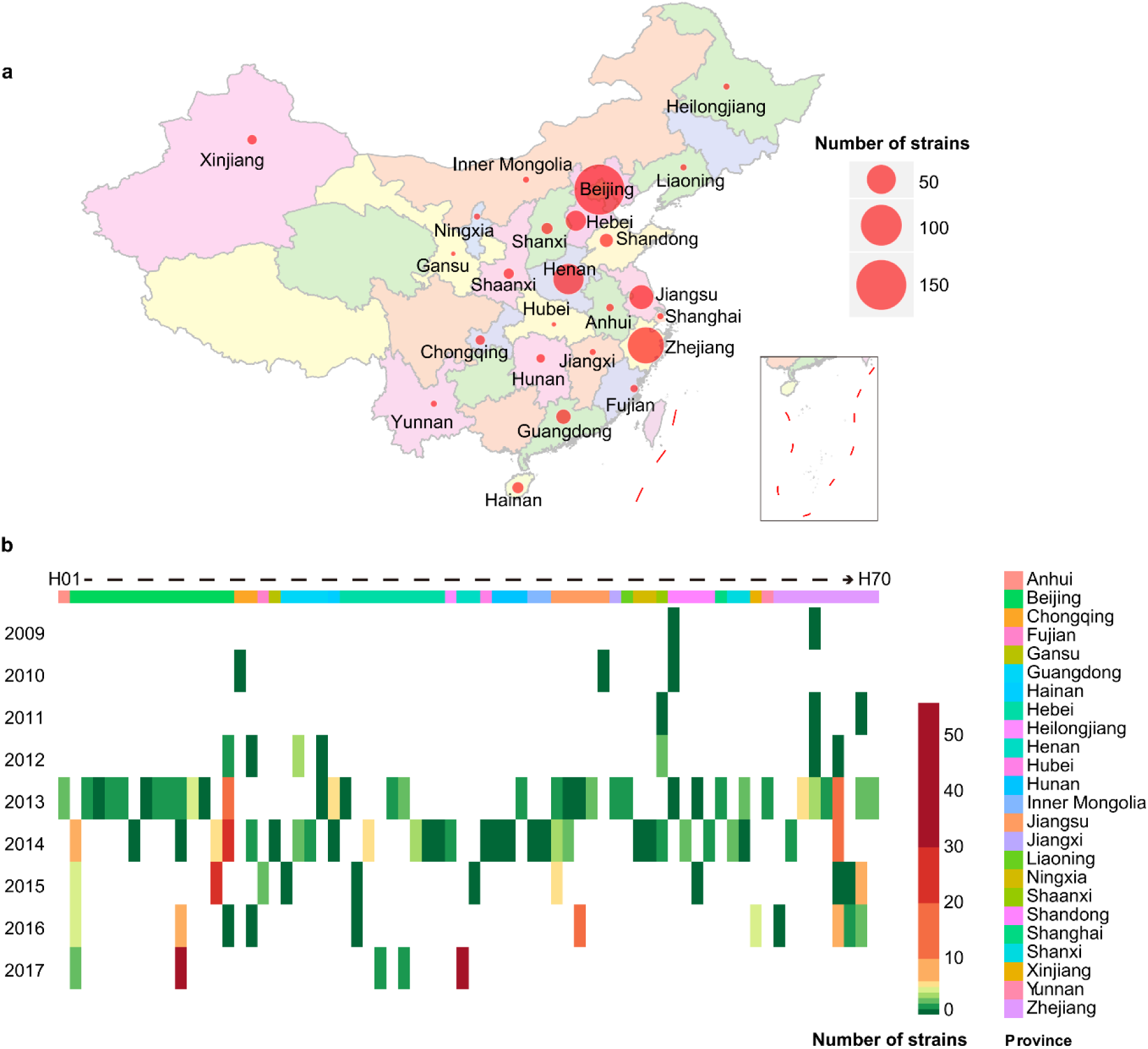
Spatial-temporal distribution of cpKP isolates from China. **a**, Distribution of our 420 cpKP isolates in different provinces. **b**, Distribution of the 420 cpKP isolates in different years and hospitals. The red circles (a) and color palettes (b) indicated the numbers of isolates. Hospital H1 to H70 (b) could be assigned to different provinces in different colors.

We first performed multilocus sequence typing (MLST; Figure 2a and Figure S1), and the results showed that 420 cpKP isolates were assigned into 48 STs, including six novel ones: ST3333, ST3334, ST3345, ST3348, ST3349 and ST3350, which can further be assigned into four singletons and 31 CGs. Among those 420 cpKp isolates, CG258 and non-CG258 account for 313 and 107, respectively. Additionally, CG258 was the most prevalent CG (313/420, 74.52%) and composed of four STs: ST11 (298/313, 95.21%), ST2667 (7/313, 2.24%), ST3348 (6/313, 1.92%) and ST258 (2/313, 0.64%). Therefore, CG258 accounts for most of the cpKP isolates, and ST11 is the overwhelmingly dominant ST of CG258 in China.

**Fig. 2.**
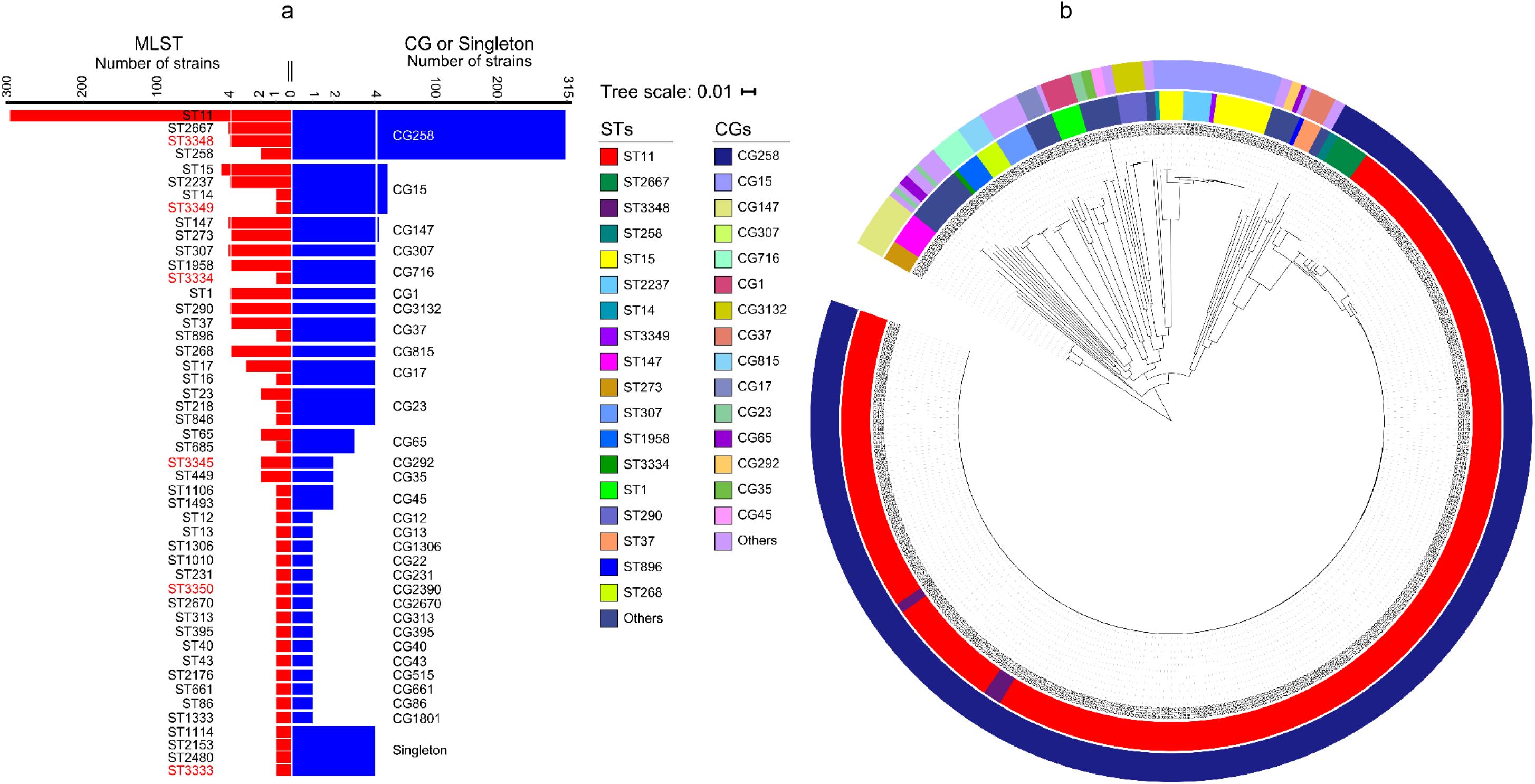
MLST and clustering tree of cpKP isolates. **a**, A profile of STs, CGs, and singletons of our 420 cpKp isolates. These 420 isolates consisted of 313 CG258 ones and 107 non-CG258 ones. The six novel STs identified in this study were highlighted in red. **b**, A maximum-likelihood clustering tree based on the 69,880 core SNPs of the 420 cpKp isolates. *K. variicola* isolate DSM 15968 was used as the outgroup but not shown in the tree. The outer and inner circles in the tree indicated CGs and STs, respectively.

We next analyzed the core-genome clustering in 2,300 global KP isolates, including our 420 cpKP isolates (Figure S2). Our 313 CG258 isolates were clustered with the other CG258 isolates, while our 107 non-CG258 isolates were scattered in the tree. Thus, in terms of genetic diversity, our 420 cpKP isolates showed good representativity.

We further performed the core-genome clustering analysis in our 420 cpKP isolates (Figure 2b). We found that the 313 CG258 isolates gathered at the farthest position from the root, suggesting CG258 was the latest clone different from other STs/CGs of cpKP. In contrast, the 107 non-CG258 isolates were located at earlier splitting branches in the tree and showed a highly dispersed pattern in the tree (containing several CGs), illustrating that they had an overall non-clonal population structure with a high level of genetic diversity.

### Phylogeny and evolutionary history of CG258

Based on the analysis of whole-genome sequencing data, the 313 CG258 cpKP isolates had a total of 6,459 core single nucleotide polymorphisms (SNPs) with an inferred *r*/*m* ratio of 3.41, indicating the presence of frequent recombination. Moreover, recombination removal generated a final collection of 233 non-redundant CG258 cpKP isolates (225 of them belonged to ST11) with 1,271 recombination-free SNPs. Based on these SNPs, a time-calibrated Bayesian maximum clade credibility (MCC) tree (Figure 3a) showed that the CG258 cpKP isolates in China could be divided into three major clades: 1 to 3. Here the Clade 1 to Clade 3 isolates had strong hospital and temporal diversity: the 23 Clade1 isolates were from 8 hospitals during 2010-2017; the 24 Clade 2 isolates were from 15 hospitals during 2011-2015; the 186 Clade3 isolates were from 42 hospitals during 2009-2017. These indicated that the formation of three major clades were not transmission events. Among the three clades, Clade 2 could be further discriminated into two subclades 2.1 and 2.2, and Clade 3 into three subclades 3.1 to 3.3. The emerging time-points of Clades 1, 2.1, 2.2, 3.1, 3.2 and 3.3 were in 1995, 2006, 2006, 2007, 2008 and 2010, respectively.

**Fig. 3.**
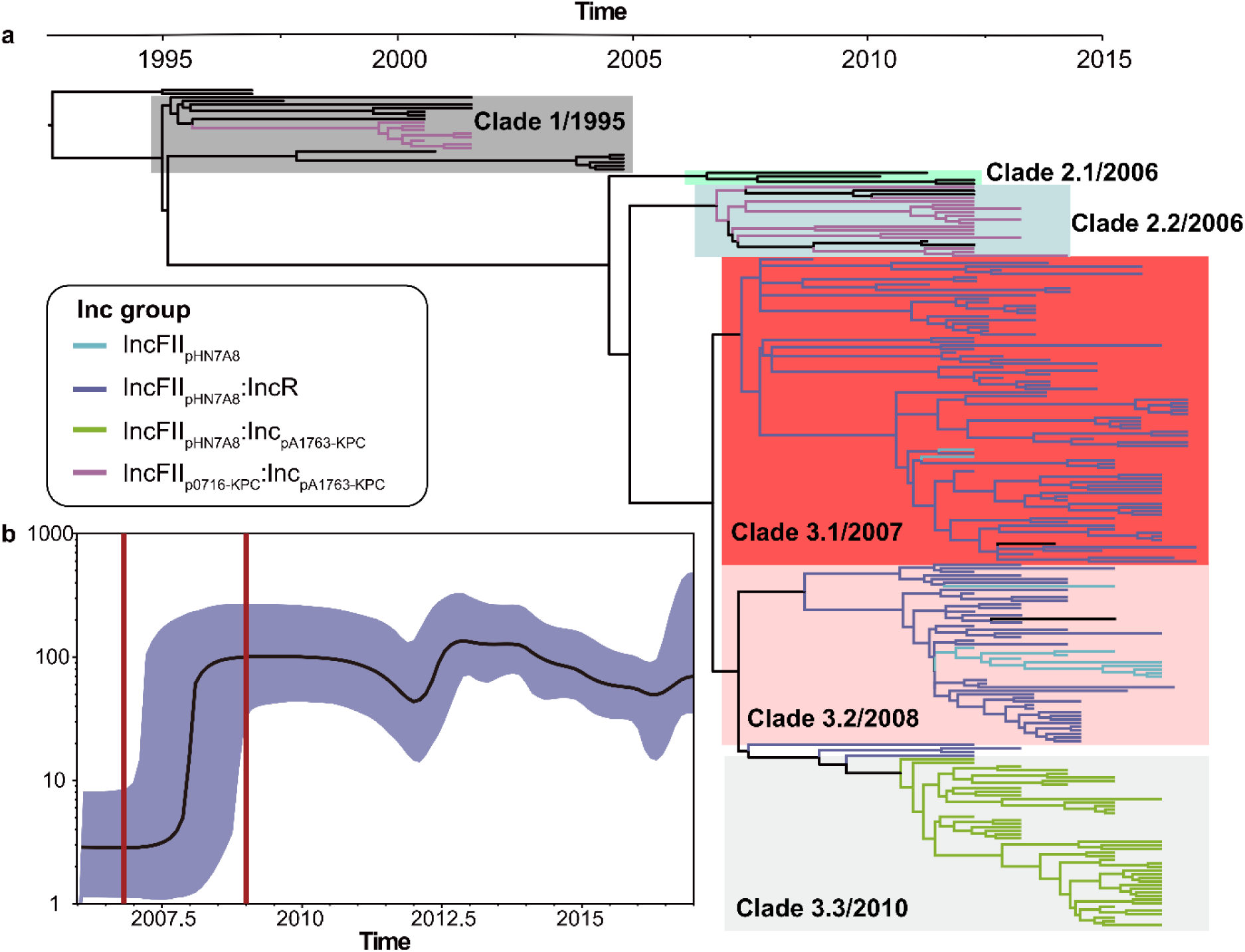
Evolutionary history of CG258 cpKP isolates. **a,** A time-calibrated MCC Bayesian phylogeny based on the 1,271 recombination-free core SNPs. These SNPs came from our 233 non-redundant CG258 cpKP isolates that were shrunk from the total collection of 313 CG258 cpKP ones (see main text). The isolates carrying different Inc groups of blaKPC-carrying plasmids were denoted as distinct colored clusters in the tree. **b,** A Bayesian skyline of the effective population size of the 233 CG258 cpKP isolates. Shadow region indicated 95% probability density interval of estimated population size.

We further performed a Bayesian skyline plot (Figure 3b) based on the above 1,271 SNPs, and the results showed a strong population expansion of CG258 cpKP during 2007-2008, which was consistent with the estimated emergence stage of Clade 3. Notably, Clade 3 accounted for 79.8% (186/233) of the CG258 cpKP isolates, and, thus, it was the dominant lineage among the three clades. Therefore, this population expansion should represent the emergence and subsequent nationwide spread of the dominant Clade 3 of CG258 cpKP in China.

### Prevalence of carbapenemase genes and *bla*_KPC_-carrying plasmids

In the 420 cpKp isolates, we identified three kinds of carbapenemase genes, including *bla*_KPC- 2/-3/-5_ (375/420, 89.29%; especially *bla*_KPC-2_ in 372 isolates), *bla*_NDM-1/-5_ (29/420, 6.9%) and *bla*_IMP-4/-38_ (19/420, 4.52%) (Table S2). Most (309/375, 82.43%) of the *bla*_KPC_-carrying isolates belong to CG258 and, meanwhile, *bla*_KPC_ was found in almost all (309/313, 98.72%) of the CG258 cpKP isolates. In contrast, the majority of the *bla*_NDM_-carrying isolates (27/29, 93.10%) and the *bla*_IMP_-carrying isolates (17/19, 89.47%) were assigned into non-CG258 (Table S3 and Figure S3). Thus, the dissemination of *bla*_KPC_, as the most prevalent carbapenemase gene, was highly associated with the spread of CG258 cpKP isolates in China.

All the detected *bla*_KPC_ genes were located in plasmids. A total of 377 *bla*_KPC_-carrying plasmids were identified from the 375 *bla*_KPC_-harboring isolates with two isolates each carrying a double copy of *bla*_KPC_ in two different plasmids. 79 of these 377 plasmids had the complete sequences (Table S1). These 377 plasmids could be assigned into 32 Inc groups, 20 (62.50%) of which had multiple replicons (Table S4). The top five Inc groups, including IncFII_pHN7A8_:IncR (n=163), IncFII_pHN7A8_:Inc_pA1763-KPC_ (n=59), IncFII_pKPHS2_:Inc_pA1763-KPC_ (n=36), IncFII_p0716-KPC_:Inc_pA1763-KPC_ (n=30) and IncFII_pHN7A8_ (n=28), accounted for 83.82% of all *bla*_KPC_-harboring plasmids (Table S4). Each of top five Inc group had at least five complete plasmid sequences (Table S1, and Figure S4 to S8). The plasmids of each Inc group carried the identical core backbone *rep* (replication) and *par* (partition) genes but showed considerable modular divergence across whole plasmid genomes (Figure S4 to S8). Overall, the *bla*_KPC_- harboring plasmids in cpKP isolates of China showed a high level of diversity, with the aforementioned top five Inc groups as the major types.

Notably, 62.50% (20/32) of Inc groups had a primary or single replicon belonging to the IncFII family, which accounted for a huge percentage of all plasmids (92.04%, 347/377) (Table S4). The IncFII replicons could be further divided into seven distinct sub-groups and displayed a high degree of genetic divergence, among which IncFII_pHN7A8_ (253/255, 99.22%) and IncFII_p0716-KPC_ (29/32, 90.63%) were highly associated with CG258. In contrast, IncFII_pKPHS2_ (35/42, 83.33%) was highly associated with non-CG258 (Figure S9).

The local *bla*_KPC_ genetic environments of the 377 *bla*_KPC_-carrying plasmids could be divided into six major types: type 1 to type 6 (Figure S10). The former five types represented different derivatives of Tn*6296* and accounted for 99.73% (376/377) of all plasmids, while type 6 was Tn*4401b* corresponding to the sole *bla*_KPC_-carrying ST258 cpKP isolate.

### Strong correlation between three major Inc groups of *bla*_KPC_-carrying plasmids and CG258

IncFII_pKPHS2_:Inc_pA1763-KPC_ and the other four top Inc groups of plasmids were highly associated with non-CG258 and CG258, respectively (Figure 4a). Firstly, 91.67% (33/36) of the *bla*_KPC_- carrying IncFII_pKPHS2_:Inc_pA1763-KPC_ plasmids came from non-CG258 involving 11 STs that could be further assigned into nine CGs (Table S3), indicating a highly dispersed dissemination of IncFII_pKPHS2_:Inc_pA1763-KPC_ plasmids among a lot of non-CG258 subtypes.

**Fig. 4.**
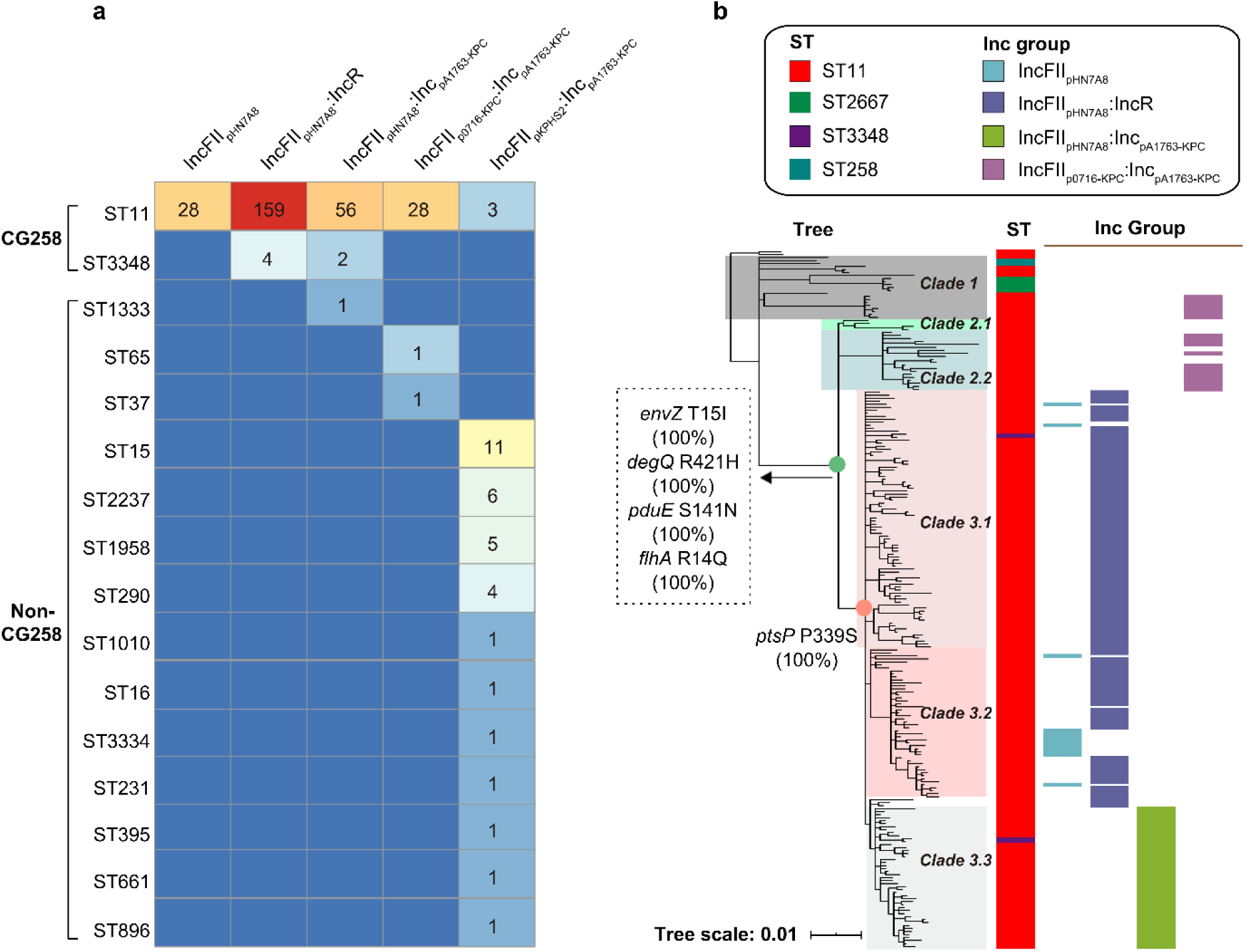
Correlation of different Inc groups of *bla*_KPC_-carrying plasmids with CG258 and non-CG258. **a,** Prevalence of the top five Inc groups (Table S4) among our CG258 and non- CG258 cpKp isolates. The numbers in the cells represented numbers of cpKp isolates (316 in total). **b**, Association of the four Inc groups with different clades of CG258. The tree was the Bayesian tree of the 233 CG258 cpKp isolates. The SNPs on the nodes indicated some specific markers for the accurate classification of Clade 1, 2, and 3 of ST11/CG258 cpKP isolates.

Secondly, 98.93% (277/280) of the other four top Inc groups of *bla*_KPC_-carrying plasmids corresponded to 88.50% of the CG258 cpKP isolates (277/313). In addition, we also downloaded 38 complete genomes of ST11/CG258 cpKP isolates from NCBI, and 77.5% ST11/CG258 cpKP isolates harbored the four Inc groups of plasmids (Table S5). Previous studies also demonstrated the strong association between ST11/CG258 cpKP and the four Inc groups of plasmids of IncFII-like plasmids (8, 21).

Moreover, different Inc groups of *bla*_KPC_-carrying plasmids had strongly correlated with various clades of CG258 (Figure 4b). We observed a strong correlation of IncFII_pHN7A8_ (13/13, 100%)+IncFII_pHN7A8_:IncR (119/122, 97.54%), IncFII_pHN7A8_:Inc_pA1763-KPC_ (45/45, 100%), and IncFII_p0716-KPC_:Inc_pA1763-KPC_ (23/23, 100%) with Clade 3.1+3.2, Clade 3.3, and Clade 2.2+an unnamed subclade of Clade 1, respectively (Figure 4b). And vice versa, Clade 3.1 +3.2 (132/136, 97.06%), Clade 3.3 (45/47, 95.74%), and Clade 2.2+an unnamed subclade of Clade 1 (23/28, 82.14%) showed a higher correlation with IncFII_pHN7A8_+IncFII_pHN7A8_:IncR, IncFII_pHN7A8_:Inc_pA1763-KPC_, and IncFII_p0716-KPC_:Inc_pA1763-KPC_, respectively (Figure 4b). In addition, we obtained four nonsynonymous and eight synonymous SNPs for discriminating Clade 2+3 from Clade 1, and one nonsynonymous and four synonymous SNPs for differentiating Clade 3 from the other two clades (Figure 4b), indicating the potential markers for accurate classification of various types of CG258 cpKP isolates.

### Acquisition of three IncFII_pHN7A8_––related Inc groups of *bla*_KPC_-carrying plasmids promoted nationwide spread of CG258

As described above, Clade 3 represented the dominant lineage of the nationwide spread CG258 cpKP in China. Almost all the Clade 3 isolates (180/186, 96.77%) contained the *bla*_KPC_-carrying plasmids of three IncFII_pHN7A8_-related Inc groups, including IncFII_pHN7A8_, IncFII_pHN7A8_:IncR, and IncFII_pHN7A8_:Inc_pA1763-KPC_ (Figure 4), which also belonged to the above top five Inc groups. This finding suggests that these three IncFII_pHN7A8_-related Inc groups had phenotypic advantages relative to the other two top Inc groups.

We next examined the susceptibility/resistance profiles of 420 cpKp isolates to nine distinct classes of 21 different antibiotics (Figure S11, and Table S1). The capability of resistance to different antibiotic classes had the following tendency (high to low): the 249 CG258 isolates harboring the *bla*_KPC_-carrying plasmids of the three IncFII_pHN7A8_-related Inc groups > the 28 CG258 isolates harboring those of IncFII_p0716-KPC_:Inc_pA1763-KPC_ > the 33 non-CG258 isolates harboring those of IncFII_pKPHS2_:Inc_pA1763-KPC_ (Figure 5a). This trend was also observed when the bacterial growth rates under antibiotic treatment were compared among the above groups of isolates (Figure 5b). In addition, we chose five representative *bla*_KPC_-carrying cpKP isolates each containing a single plasmid: G134 (ST11+IncFII_pHN7A8_), G285 (ST11+IncFII_pHN7A8_:IncR), G318 (ST11+IncFII_pHN7A8_:Inc_pA1763-KPC_), G165 (ST11+IncFII_p0716-KPC_:Inc_pA1763-KPC_) and G344 (non-CG258+IncFII_pKPHS2_:Inc_pA1763-KPC_) for *in vitro* competitive assay. The results showed a similar tendency: the three IncFII_pHN7A8_-related Inc groups > IncFII_p0716-KPC_:Inc_pA1763-KPC_ > IncFII_pKPHS2_:Inc_pA1763-KPC_ (Figure 5c). Therefore, in relative to IncFII_p0716-KPC_:Inc_pA1763-KPC_ and IncFII_pKPHS2_:Inc_pA1763-KPC_, the acquisition of three IncFII_pHN7A8_-related Inc groups of *bla*_KPC_- carrying plasmids rendered their host KP higher levels of antibiotic resistance, growth, and competition advantages. These phenotypic advantages might promote the nationwide spread of the dominant Clade 3 of CG258 cpKP in China.

**Fig. 5.**
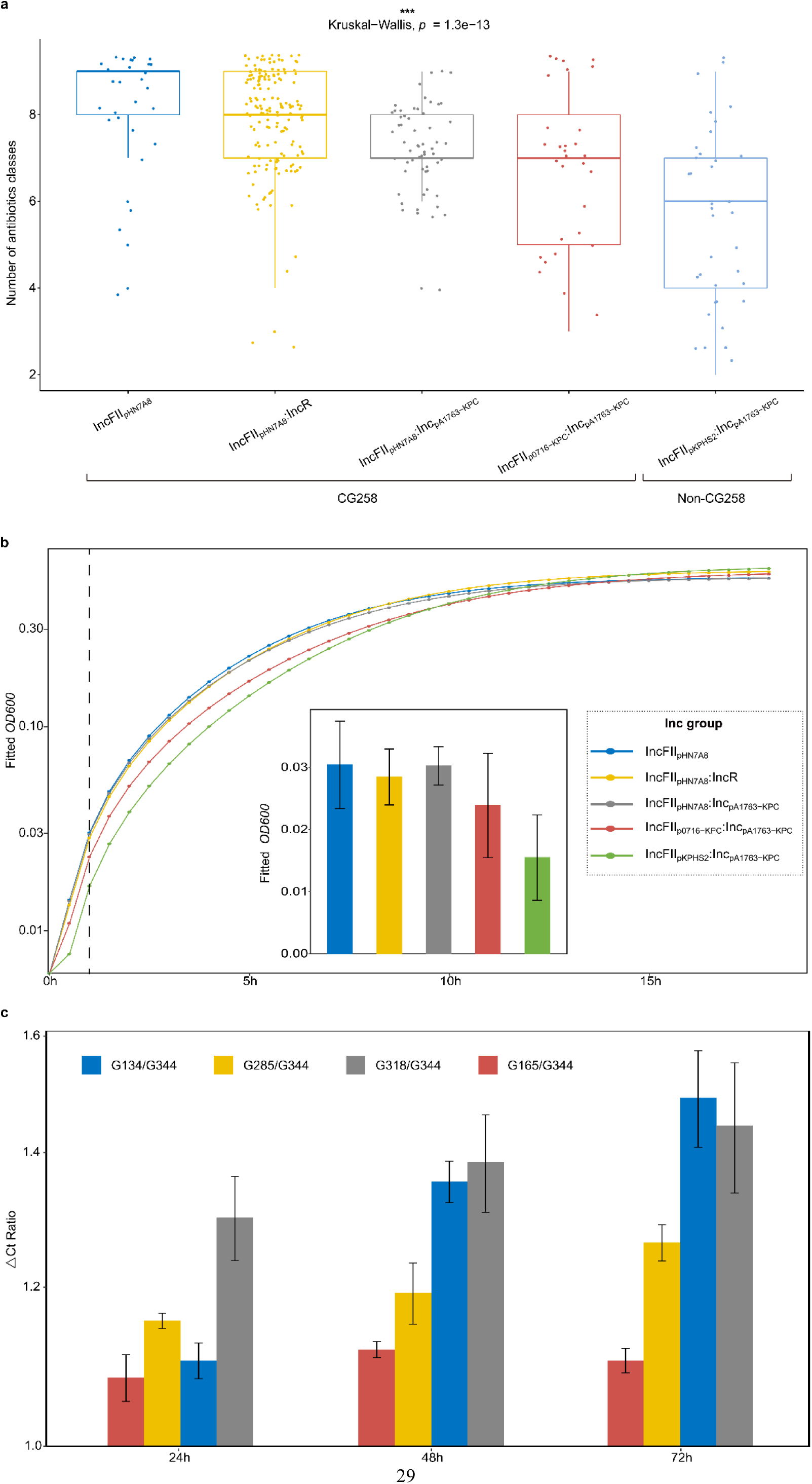
Resistance, growth, and competition advantages of cpKP harboring *bla*_KPC_- carrying plasmids of different Inc groups. **a,** Boxplots showed the numbers of classes of antibiotics that different subgroups of cpKP isolates were resistant to. The *p* values were obtained using Kruskal-Wallis, a non-parametric test for the comparison of multiple groups. ***, statistically significant with *p* < 0.0001. **b,** Bacterial growth curves of different subgroups of cpKP isolates. The dashed line indicated the timepoint at 1 h, when bacteria were at the logarithmic growth phases; corresponding *OD_600_* values (mean ± standard error) were shown in the embedded bar-plot. **c,** Bacterial *in vitro* competition experiments. Shown were the Ct value ratios (mean ± standard error) between each two cpKP isolates at 24h, 48h, and 72h, respectively.

### Optimized antibiotic combination regimens for cpKP treatment

All of the cpKP isolates were resistant to β-lactams including carbapenems, whereas the resistant rates of these isolates to aminoglycosides [amikacin (47.86%, 201/420), tobramycin (57.62%, 242/420) and gentamicin (79.52%, 334/420)] and trimethoprim/sulfamethoxazole (65%, 273/420) were much lower (Figure S11a). For each of the ten antibiotics tested (Figure S11b), CG258 isolates displayed a much higher drug resistance rate than non-CG258. As shown by the number of antibiotic classes that the isolates were resistant to, CG258 showed much higher resistance levels than non-CG258 (Figure S12). Specifically, CG258 had the lowest resistance rates for amikacin (57.4%, 179/312) followed by trimethoprim/sulfamethoxazole (67.7%, 212/313) and tobramycin (73.2%, 202/276), while non-CG258 exhibited the lowest resistance for amikacin (20.6%, 22/107) followed by tobramycin (50.6%, 40/79) and levofloxacin (53.9%, 55/102) (Figure S11b). This large discrepancy of resistance profile between CG258 and non-CG258 led us to optimize the antibiotic combination regimens to treat CG258 and non-CG258.

Based on the calculated resistance ratios when different two-antibiotics combination regimens were used for CG258 treatment, two optimized combinations of ‘amikacin+trimethoprim/sulfamethoxazole’ and ‘tobramycin+trimethoprim/sulfamethoxazole’ produced the resistance ratios of 32.59% (102/313) and 39.62% (124/313), respectively, which were much lower than the ratio of 57.4% (179/312) calculated for single-antibiotic ‘amikacin’ (Figure 6a). In addition, the combinations of ‘amikacin+macrodantin’ and ‘amikacin+cefotetan’ represented the two optimized two-antibiotics combination for non-CG258 treatment, with the resistance ratios of 13.08% (14/107) and 14.02% (15/107), respectively (Figure 6b).

**Fig. 6.**
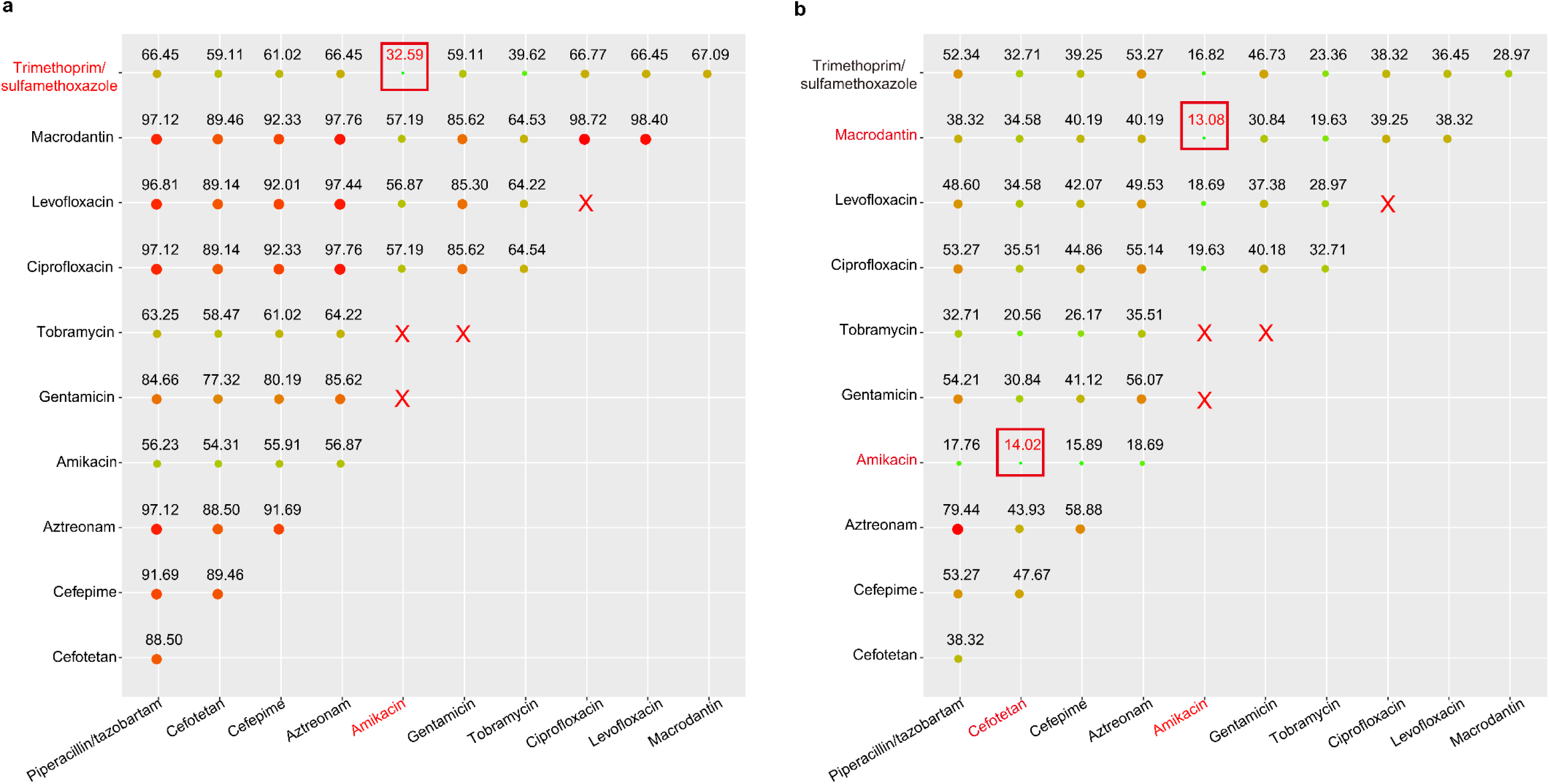
Optimized two-antibiotics combination regimens for treatment of CG258 and non-CG258. Shown were the resistance ratios (the numbers of isolates resistant to both antibiotics tested/the total numbers of isolates) when different two-antibiotics combination regimens were used for the treatment of CG258 (a) and non-CG258 (b). The drug resistance ratio of each two different drug combinations of 11 antibiotics (excluding 10 antibiotics with the resistance ratio of ∼100% and some banned drug combinations) was calculated. The size of the solid circles increases with the increase in resistance ratio. The red rectangle indicates the optimized drug combinations of two drugs. The red cross indicates the prohibited drug combinations since they belong to the same type of antimicrobial drug.

## Discussion

This study primarily revealed that the three host-plasmid ‘golden’ combinations played an important and even decisive role in the successfully clonal spread and dissemination of ST11/CG258 cpKp in China. Here the three host-plasmid ‘golden’ combinations indicate the three main phylogenetic subclades of ST11/CG258 cpKp isolates carrying the three most prevalent Inc groups of KPC-producing plasmids: Clade 3.1+Clade 3.2 — IncFII_pHN7A8_, Clade 3.1+Clade 3.2 —IncFII_pHN7A8_:IncR, Clade 3.3 — IncFII_pHN7A8_:Inc_pA1763-KPC_ (Figure 3b and Figure 4b). We name them “golden combinations” due to the genotypic and phenotypic advantages of these isolates. On the one hand, our findings illustrated the genotypic advantages of the three host-plasmid ‘golden’ combinations in strong correlation and coevolution (Figure 3a and Figure 4b); on the other hand, we demonstrated the phenotypic advantages of the three ‘golden’ combinations in drug-resistance, growth and competition (Figure 5). The genotypic and phenotypic advantages of the three host-plasmid ‘golden’ combinations are reciprocal causation, which improve the adaptability of the three IncFII_pHN7A8_ families of plasmids to ST11/CG258 cpKp isolates, further leading to their successfully clonal spread and dissemination in China. Overall, they form a closed-loop process: the genotypic and phenotypic advantages of three host-plasmid ‘golden’ combinations, and their successful spread and dissemination in China (Figure 7).

**Fig. 7.**
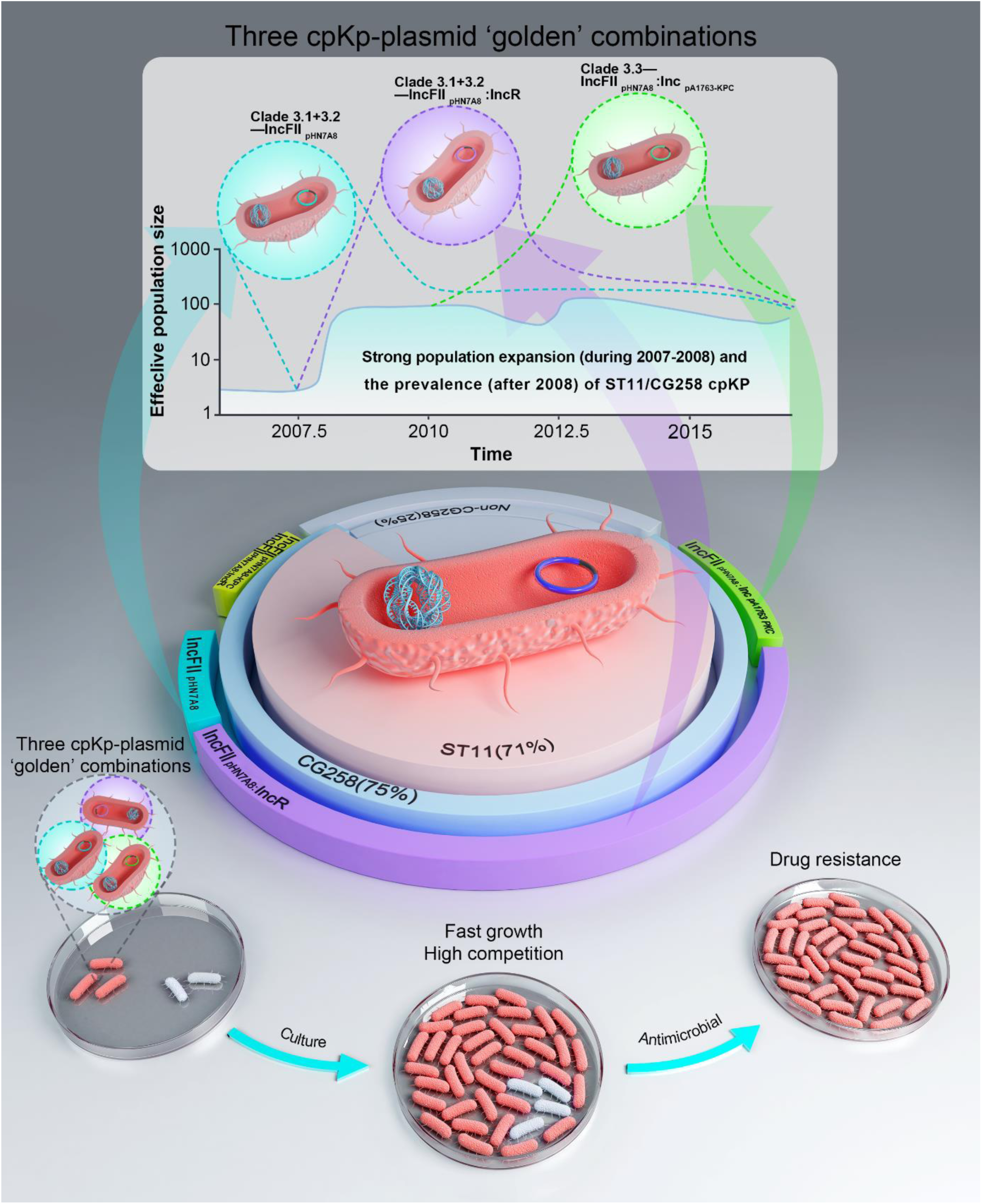
Three ‘golden’ host––plasmid combinations. The three ‘golden’ host––plasmid combinations (Clade 3.1+3.2—IncFII_pHN7A8_, Clade 3.1+3.2—IncFII_pHN7A8_:IncR, Clade 3.3— IncFII_pHN7A8_:Inc_pA1763-KPC_) led to the strong population expansion during 2007-2008 and subsequent maintenance of the prevalence and clonal dissemination of ST11/CG258 cpKP after 2008. They endowed cpKP with the phenotypes advantages in growth, competition and resistance.

### Three ‘golden’ combinations of host KP––*bla*_KPC_-carrying plasmid constitutes the major genetic basis (main internal cause) for the nationwide spread of ST11/CG258 cpKP in China

Most cpKP isolates in China belong to the clonal group CG258 with ST11 as the dominant ST. In contrast, non-CG258 isolates had an overall non-clonal population structure with very high levels of genetic diversity. Our core-genome phylogenetic analysis identified Clade 3 as the dominant lineage. On the other hand, *bla*_KPC_ was the dominant carbapenemase gene located in the plasmids of cpKP isolates. Among the five major Inc groups of *bla*_KPC_-harboring plasmids, a strong correlation of IncFII_pHN7A8_+IncFII_pHN7A8_:IncR, IncFII_pHN7A8_:Inc_pA1763-KPC_, and IncFII_p0716-KPC_:Inc_pA1763-KPC_ with CG258 Clade 3.1+3.2, Clade 3.3, and Clade 2.2+an unnamed subclade of Clade 1, respectively, were observed, while IncFII_pKPHS2_:Inc_pA1763-KPC_ was highly associated with non-CG258.

The dissemination of CG258 cpKP in China was characteristic of a nationwide spread of the dominant Clade 3 since 2007-2008 (Figure 3b). A strong population expansion was observed for CG258 cpKP during 2007-2008, which was accompanied by the emergence of two host-plasmid ‘golden’ combinations at the same period: Clade 3.1+Clade 3.2 — IncFII_pHN7A8_, Clade 3.1+Clade 3.2 —IncFII_pHN7A8_:IncR. In addition, the host-plasmid ‘golden’ combination, Clade 3.3 — IncFII_pHN7A8_:Inc_pA1763-KPC_ emerged at 2010. The expanded CG258 population retained at a high level with limited dynamic fluctuation, which was linked to the spread of Clade 3.1+3.2 since 2007-2008 as well as that of Clade 3.3 since 2010 (Figure 3b and Figure 7).

Acquisition of *bla*_KPC_-carrying plasmids of IncFII_pHN7A8_, IncFII_pHN7A8_:IncR, and IncFII_pHN7A8_:Inc_pA1763-KPC_ endowed their host KP isolates with phenotypes of a higher competitive growth rate and more antibiotic resistance, relative to those harboring IncFII_p0716- KPC_:Inc_pA1763-KPC_ or IncFII_pKPHS2_:Inc_pA1763-KPC_ (Figure 5). Remarkably, the evolution of three ‘golden’ combinations of host CG258––*bla*_KPC_-carrying plasmid (Clade 3.1+3.2––IncFII_pHN7A8_, Clade 3.1+3.2––IncFII_pHN7A8_:IncR, and Clade 3.3––IncFII_pHN7A8_:Inc_pA1763-KPC_) rendered them phenotypic advantages that might directly lead to their strong expansion and subsequent clonal dissemination since 2007-2008 in China.

In particular, more attention should be paid to the following two ‘golden’ combinations: Clade 3.1+3.2––IncFII_pHN7A8_:IncR, and Clade 3.3––IncFII_pHN7A8_:Inc_pA1763-KPC_, as they were ranked at the first (58.84%, 163/277) and second (20.94%, 58/277) percentages of CG258 cpKP isolates, respectively, and positioned at the farthest point from the root of the phylogenetic tree of CG258 cpKP isolates. Thus, they might represent the latest differentiation events and have the best host KP––plasmid adaptability, therefore, emerge as the highest risk lineages.

IncFII_pHN7A8_:IncR or IncFII_pHN7A8_:Inc_pA1763-KPC_ plasmids are the chimeras of IncFII_pHN7A8_ plasmids (carrying *bla*_KPC_) and IncR/Inc_pA1763-KPC_ plasmids (carrying multiple resistance genes) (22–25). Compared to single-replicon plasmids, multi-replicon plasmids have the advantages of rapid replication, replicon substitution, multiple partitioning and/or toxin-antitoxin systems for plasmid maintenance, and a high survival rate. The above two chimeric plasmids also possess additional unique advantages: i) IncR or Inc_pA1763-KPC_ backbones are very small, giving the low adaptability costs of the chimeras after their fusion with IncFII_pHN7A8_; ii) IncR or Inc_pA1763-KPC_ plasmids carried large multidrug resistance regions, expanding the resistance profile of their hosts; and iii) although IncR or Inc_pA1763-KPC_ plasmids did not carry conjugation transfer regions, they can obtain self-transfer ability after fusion with conjugative IncFII_pHN7A8_ plasmids, further facilitating the spread of resistance genes they carried.

### Widespread use of carbapenems is the leading external cause for the expansion and prevalence of ST11/CG258 cpKp in China

Bacterial antibiotic resistance as a serious threat to public health is of the greatest concern in China. In addition to the genetic basis of three ‘golden’ combinations accounting for the population size change of CG258 cpKP, rapidly increased sales/use of carbapenems since 2005 with the highest growth rate during 2007-2008 (Figure S13) constitutes the major external force driving antibiotic resistance, clonal expansion and consequently nationwide spread of ST11/CG258 cpKP in China.

Specifically, the consumption of Meropenem (a common carbapenem antibiotic) increased sharply from 2007 to 2009 (Figure S13). At the same period, some other carbapenems were listed in China and also facilitated their consumption (ertapenem: 2005, faropenem: 2006, biapenem: 2008). Overall, the rapid growth of carbapenem consumption around 2007 might directly led to the expansion of ST11/CG258 cpKP population. After 2007-08, the expansion of cpKP population was restrained and entered a plateau (Figure 3a), which was mainly due to the slow growth of sales of carbapenems in China. Along with imposing restrictions on antibiotic prescription issued by the Ministry of Health of China since 2011, it has been reported that the use of antibiotics in many hospitals across the country has decreased significantly (26, 27). In addition, the Chinese government has introduced several policies to strictly control the usage of carbapenem antibiotics covering 1,429 hospitals (http://www.chinets.com).

In conclusion, the population change of ST11/CG258 cpKp isolates in China might be derived from the internal cause (three host-plasmid ‘golden’ combinations) and the external cause (widespread use of carbapenems). Exogenous variables could result in endogenous changes, and they worked together to facilitated the spread and prevalence of ST11/CG258 cpKP isolates in China. Internal monitoring and external control should effectively inhibit the epidemic spread of cpKP isolates.

### Possibility of precision medication in clinic cpKP treatment

Based on the susceptibility/resistance phenotypes to nine distinct classes of 21 different antibiotics, cpKP isolates in China displayed the lowest resistance rates for three aminoglycosides (amikacin, tobramycin, gentamicin) and trimethoprim/sulfamethoxazole. Similar results have also been reported in a wealth of literature (Table S6). Additionally, our findings revealed that CG258 cpKP has much broader resistance profiles than non-CG258 cpKP (Figure S12), which is supported by the epidemiological survey data of carbapenem-resistant *Enterobacteriaceae* from 2012 to 2016 in China (19). These findings make the possibility to optimize the two-antibiotics combination regimens for treatment of CG258 and non-CG258 through calculating their resistance ratios when different antibiotic combinations are used. More importantly, cpKP with three ‘golden’ combinations showed the highest resistance profiles than other cpKP. More in-depth studies on different antibiotic combination regimens need to be executed for various cpKP genotypes with different Inc groups of plasmids, which will provide a precise reference and a choice to treat cpKP infection by using the existing drugs effectively. Eventually, we hope that our study can appeal for people to closely monitor the adaptability of resistant plasmids, so as to effectively prevent the emergence/dissemination of new ‘golden’ host-plasmid combinations. Additionally, we should consider both genotypes (isolates) and Inc groups (drug-resistant plasmids) to achieve precision medication of cpKp infections.

## Materials and Methods

### Bacterial isolates and genomic DNA extraction

We collected 2,803 clinical *K. pneumoniae* isolates from 70 hospitals in 24 provinces of China from 2009 to 2017. After eliminating 59 culture failed isolates, 2,744 isolates were obtained for PCR detection of *K. pneumoniae*-specific *khe* gene to identify the species. After excluding the 53 isolates without *khe* gene, 2,691 isoates were further tested to produce carbapenemases by Modified Carba NP test: 493 isolates were confirmed to produce carbapenemases. Bacterial genomic DNAs were then extracted using a Qiagen UltraClean Microbial DNA Isolation Kit, which were sequenced by Illumina technology. After excluding 73 low-quality sequencing sample, 420 cpKP genomes were used for the subsequent analysis (Table S1).

### Genome sequencing and assembly

The draft genome sequences of bacterial genomic DNA were sequenced from a paired-end library with an average insert size of 350 bp on an Illumina HiSeq2000 sequencing platform (28). Adapters and low-quality reads were removed using *FASTX-Toolkit* (http://hannonlab.cshl.edu/fastx_toolkit/). *SPAdes* v3.9.0 (29) was used to do the *de novo* assembly from the trimmed sequence reads using *k*-mer sizes of 21, 33, 55 and the -cov-cutoff flag set to ‘auto’. Isolates were discarded through the following analyses: i) if the size of the *de novo* assembly was outside of 5-7 Mb, ii) if the average nucleotide identity to NJST258_1 was lower than 95% or the top match was not NJST258_1 after the *de novo* assembly was compared with the reference genomes of five *Klebsiella* spp. (*K. pneumoniae* NJST258_1, CP006923; *K. quasipneumoniae* ATCC 700603, CP014696; *K. michiganensis* E718, NC_018106; *K. oxytoca* CAV1374, CP011636; *K. variicola* DSM 15968, CP010523) using *Pyani-0.2.7* (30), and iii) if the percentage of the total number of genomic sites with more than 10-fold depth of coverage was lower than 80% after the raw sequencing reads of each isolate were mapped to the NJST258_1 genome using *Bowtie2* (31) and the depth of coverage for each position on the genome were calculated using *samtools depth* v0.1.19 (32).

The complete genome sequences was obtained from a sheared DNA library with an average size of 10 kb on a PacBio RSII sequencer (Pacific Biosciences) (33), and the *de novo* sequence assembly was performed using *SMRT Analysis* v2.3.0 (https://smrt-analysis.readthedocs.io/en/latest/). Nanopore GridION platform was also used for whole-genome sequencing (34), and the high-quality reads (mean_qscore_template ≥7 and length ≥ 1,000) were screened for further *de novo* sequence assembly using *canu* (https://canu.readthedocs.io/en/latest/) (35). Circularization of chromosomal or plasmid sequences was achieved by manual comparison. *Pilon* v.1.13 (36) was employed to polish complete genome sequences using Illumina sequencing reads.

### MLST

*SRST2* (37) was used to identify the ST of each KP isolate by mapping its Illumina sequencing reads to the Pasteur *Klebsiella* MLST Database (http://bigsdb.pasteur.fr/klebsiella/klebsiella.html). All the STs in the *Klebsiella* MLST database (last accessed August 3, 2018) were assigned to different CGs using *eBURST* (38).

### Construction of maximum-likelihood clustering trees

Chromosome sequences were mapped to a reference sequence of NJST258_1 (11) using *Bowtie2* (31). The core SNPs of the 2,300 global KP isolates were identified using *Mummer* v3.25 (39), and the core SNPs of our 420 cpKp isolates were called using *GATK Unified Genotyper* (40). We filtered all the SNPs in the repetitive DNA regions (identified by *RepeatMasker*, http://www.repeatmasker.org/) and the mobile genetic elements (including insertion sequences, transposons, integrons, and phage-related genes). Based on the above core SNPs, the maximum-likelihood clustering trees of the 2,300 global KP isolates and our 420 cpKp isolates were constructed using *FastTree* V2.1.9 (41) and *RAxML* (42), respectively (Bootstrap value: 500).

### Construction of recombination-free Bayesian phylogenetic tree

Our 313 CG258 cpKP isolates were subjective to sequence alignment. Recombination DNA regions were predicted using *ClonalFrameML* (43), followed by removal of all putative recombinant SNP sites. A Bayesian phylogenetic tree was constructed from the recombination-free core SNPs of the resulting 233 non-redundant CG258 cpKP isolates using *MrBayes* (44) and visualized using *iTOL* (https://itol.embl.de/).

### Bayesian phylogenetic inference and molecular dating analyses

Bayesian skyline analysis was performed to calculate the change in the effective population size of the above 233 isolates using *BEAST* v1.8.4 (45). The three standard substitution models, Hasegawa-Kishino-Yano (HKY), general time-reversible (GTR), and Tamura-Nei 93 (TN93) was tested in combination with the estimated/empirical base frequency, the gamma (G) site heterogeneity and the loose molecular clock. By testing various parameter combinations, the model combination GTR+empirical+G4 was selected. The tip date was defined as the sampling time. In the end, three independent chains of 5×10^7^ generations were run to ensure calculation accuracy, with sampling every 1,000 iterations. The resulting Bayesian skyline plot was visualized using *Tracer* v1.7 (46). A time-calibrated Bayesian MCC tree of the above 233 isolates was constructed using *TreeAnnotator* (https://beast.community/treeannotator) and visualized using *FigTree* (http://tree.bio.ed.ac.uk/software/figtree/).

### Plasmid analysis

All the fully sequenced *bla*_KPC_-carrying plasmids from GenBank (last accessed Aug 29, 2018) and our study were used as the references. The draft sequences of the rest *bla*_KPC_-carrying plasmids in our 420 cpKp isolates were aligned using *BLAST* (47) and custom *Perl* scripts. Inc groups and core backbone *rep* and *par* genes were determined for all the *bla*_KPC_-carrying plasmids in our 420 cpKp isolates. To ensure accuracy, the assembled draft plasmid sequences met the following three criteria (48): the *bla*_KPC_-embedded contigs had 100% query coverage and ≥ 99% identity with corresponding reference plasmids; the *bla*_KPC_-embedded contigs and the *rep*-embedded contigs of the same plasmid had similar sequencing depths; each draft plasmid sequence had ≥ 70% Query coverage and ≥ 94% identity with corresponding reference plasmids.

### Identification of carbapenemase genes

The major plasmid-borne carbapenemase genes were screened for each cpKP isolate by PCR, followed by amplicon sequencing using ABI 3730 Sequencer (49). The variants of *bla*_KPC_, *bla*_NDM_, and *bla*_IMP_ were identified from genome sequence data using *ResFinder* (50).

### Bacterial phenotypic resistance assays

Bacterial antimicrobial susceptibility was tested by BioMérieux VITEK 2 and interpreted based on the 2018 Clinical and Laboratory Standards Institute (CLSI) guidelines (51). The activity of Ambler class A/B/D carbapenemases in bacterial cell extracts was determined by a modified CarbaNP test (49).

### Bacterial growth curves

Bacterial growth curves were measured on a 96 well-microtitre plate using a Thermo Scientific Multiskan FC instrument. Equivalent amount of overnight bacterial culture was inoculated in each well containing 200 μl of LB liquid medium (4 mg/L meropenem), and the mixtures were cultured at 37°C overnight with a speed of 5 Hz. The bacterial growth curve was determined through a course of time by recording the turbidity at 600 nm using the microplate reader of the Multiskan FC instrument. Experiments were performed in triplicate.

### *In vitro* competition experiments

Equivalent amount of overnight bacterial cultures of two indicated bacterial isolates were inoculated into 10 ml of LB liquid medium (4 mg/L meropenem), and the mixtures were cultured at 37°C for 72 h in a shaker with a speed of 200 rpm. At 0, 24, 48, and 72h, 3 mL aliquots of the cultures were taken, and genomic DNAs were extracted. To examine the competition between two bacterial isolates, real-time qPCRs were performed to determine the ratio of Ct values between each of the four ST11/CG258 isolates (G134, G285, G318, and G165) and the control non-CG258 isolate G344. The five genes G134_05212, G285_01367, G318_02254, G165_02217, and G344_00764 were selected as PCR target sequences, and the corresponding PCR primers were listed in Table S7. Experiments were performed in triplicate.

### Data availability

The genome sequences in this study were submitted to Genome Sequence Archive under accession number CRA003059. Individual accession numbers for sequence data were also available in Table S1.

## Author Contributions

DZ and FC conceived the study. CL and TY performed the bioinformatics analyses. XJ, YJ, LY, GM, ZY, YJ, XL, and XW carried out the experimental analyses. CL, XJ, TY, XY, and SL drew the figures. DZ, FC, and CL wrote the manuscript. All authors read and approved the final manuscript.

## Supplementary Information

**Table S1 presented as a separate document**

**Table S2.** Carbapenemase genes from our 420 cpKP isolates.

| **Carbapenemase gene** | **Number of isolates (%)** |
| --- | --- |
| *bla*<sub>KPC</sub> | 372 (88.57) |
| *bla*<sub>KPC-2</sub> | 369 (87.86) |
| *bla*<sub>KPC-3</sub> | 1 (0.24) |
| *bla*<sub>KPC-5</sub> | 2 (0.48) |
| *bla*<sub>NDM</sub> | 26 (6.19) |
| *bla*<sub>NDM-1</sub> | 25 (5.95) |
| *bla*<sub>NDM-5</sub> | 1 (0.24) |
| *bla*<sub>IMP</sub> | 19 (4.52) |
| *bla*<sub>IMP-4</sub> | 13 (3.10) |
| *bla*<sub>IMP-38</sub> | 6 (1.43) |
| *bla*<sub>KPC</sub>+*bla*<sub>NDM</sub> | 3 (0.71) |
| *bla*<sub>KPC-2</sub>+*bla*<sub>NDM-1</sub> | 3 (0.71) |

**Table S3.** Carbapenemase genes from different STs/CGs of our 420 cpKp isolates.

| CG | ST | Number of isolates (%) |  |  |  |  |
| --- | --- | --- | --- | --- | --- | --- |
|  |  | Total | *bla*KPC -<br>carrying | *bla*NDM -<br>carrying | *bla*KPC- and<br>*bla*NDM -<br>carrying | *bla*IMP -<br>carrying |
| CG258 | ST11 | 298 | 295 (99) |  | 1 (0.3) | 2 (0.7) |
|  | ST2667 | 7 | 7 (100) |  |  |  |
|  | ST3348 | 6 | 6 (100) |  |  |  |
|  | ST258 | 2 | 1 (50.0) | 1 (50.0) |  |  |
| CG1 | ST1 | 6 | 2 (33.3) | 4 (66.7) |  |  |
| CG12 | ST12 | 1 | 1 (100) |  |  |  |
| CG13 | ST13 | 1 | 1 (100) |  |  |  |
| CG15 | ST15 | 17 | 16 (94.1) |  |  | 1 (5.9) |
|  | ST2237 | 6 | 6 (100) |  |  |  |
|  | ST3349 | 1 |  | 1 (100) |  |  |
|  | ST14 | 1 |  |  |  | 1 (100) |
| CG17 | ST17 | 3 |  | 1 (33.3) |  | 2 (66.7) |
|  | ST16 | 1 | 1 (100) |  |  |  |
| CG22 | ST1010 | 1 | 1 (100) |  |  |  |
| CG23 | ST23 | 2 | 2 (100) |  |  |  |
|  | ST846 | 1 |  |  |  | 1 (100) |
|  | ST218 | 1 | 1 (100) |  |  |  |
| CG35 | ST449 | 2 |  |  | 2 (100) |  |
| CG37 | ST37 | 4 | 2 (50.0) | 1 (25.0) |  | 1 (25.0) |
|  | ST896 | 1 | 1 (100) |  |  |  |
| CG40 | ST40 | 1 |  | 1 (100) |  |  |
| CG43 | ST43 | 1 |  |  |  | 1 (100) |
| CG45 | ST1493 | 1 |  | 1 (100) |  |  |
|  | ST1106 | 1 | 1 (100) |  |  |  |
| CG65 | ST65 | 2 | 2 (100) |  |  |  |
|  | ST685 | 1 | 1 (100) |  |  |  |
| CG86 | ST86 | 1 | 1 (100) |  |  |  |
| CG147 | ST147 | 7 | 2 (28.6) | 5 (71.4) |  |  |
|  | ST273 | 4 |  | 4 (100) |  |  |
| CG231 | ST231 | 1 | 1 (100) |  |  |  |
| CG292 | ST3345 | 2 |  |  |  | 2 (100) |
| CG307 | ST307 | 7 | 1 (14.3) |  |  | 6 (85.7) |
| CG313 | ST313 | 1 |  | 1 (100) |  |  |
| CG395 | ST395 | 1 | 1 (100) |  |  |  |
| CG515 | ST2176 | 1 |  |  |  | 1 (100) |
| CG661 | ST661 | 1 | 1 (100) |  |  |  |
| CG716 | ST1958 | 5 | 5 (100) |  |  |  |
|  | ST3334 | 1 | 1 (100) |  |  |  |
| CG815 | ST268 | 5 | 5 (100) |  |  |  |
| CG1306 | ST1306 | 1 |  | 1 (100) |  |  |
| CG2390 | ST3350 | 1 |  | 1 (100) |  |  |
| CG2670 | ST2670 | 1 |  | 1 (100) |  |  |
| CG3132 | ST290 | 6 | 5 (83.3) | 1 (16.7) |  |  |
| CG1801 | ST1333 | 1 | 1 (100) |  |  |  |
| NA | ST2480 | 1 |  | 1 (100) |  |  |
| NA | ST3333 | 1 |  | 1 (100) |  |  |
| NA | ST1114 | 1 |  |  |  | 1 (100) |
| NA | ST2153 | 1 | 1 (100) |  |  |  |

**Table S4.** Inc groups of the 377 bla_KPC_-carrying plasmids*.

| **Inc group**<br>**(n=32)** | **Number of plasmids**<br>**(n=377)** | **%Percent** |
| --- | --- | --- |
| IncFII<sub>pHN7A8</sub> | 28 | 7.43 |
| IncFII<sub>pHN7A8</sub>:IncR | 163 | 43.24 |
| IncFII<sub>pHN7A8</sub>:Inc<sub>pA1763</sub>-KPC | 59 | 15.65 |
| IncFII<sub>pHN7A8</sub>:IncN1 | 1 | 0.27 |
| IncFII<sub>pHN7A8</sub>:IncN1:IncR | 2 | 0.53 |
| IncFII<sub>pHN7A8</sub>:Inc<sub>pA1763</sub>-KPC:IncN1 | 2 | 0.53 |
| IncFII<sub>pHN7A8</sub>:IncFIB<sub>pFB2.3</sub> | 1 | 0.27 |
| IncFII<sub>pKPHS2</sub>:Inc<sub>pA1763</sub>-KPC | 36 | 9.55 |
| IncFII<sub>pKPHS2</sub>:IncR | 5 | 1.33 |
| IncFII<sub>pKPHS2</sub>:IncFIB<sub>pSC138</sub> | 1 | 0.27 |
| IncFII<sub>pKPHS2</sub>:Inc<sub>pLT968725</sub> | 1 | 0.27 |
| IncFII<sub>pKPHS2</sub>:Inc<sub>pA1763</sub>-KPC:IncR | 2 | 0.53 |
| IncFII<sub>p0716</sub>-KPC:Inc<sub>pA1763</sub>-KPC | 30 | 7.96 |
| IncFII<sub>p0716</sub>-KPC:IncR | 1 | 0.27 |
| IncFII<sub>p0716</sub>-KPC:IncFIB<sub>pSC138</sub> | 1 | 0.27 |
| IncFII<sub>pCP020359</sub>:Inc<sub>pA1763</sub>-KPC | 7 | 1.86 |
| IncFII<sub>R100</sub> | 4 | 1.06 |
| IncFII<sub>R100</sub>:IncFIB<sub>plasmid F</sub> | 1 | 0.27 |
| IncFII<sub>R100</sub>:IncFIA<sub>pBK30661</sub> | 1 | 0.27 |
| IncFII<sub>pBK30683</sub> | 1 | 0.27 |
| IncX6 | 10 | 2.65 |
| IncR | 3 | 0.80 |
| IncR:IncFIB<sub>plasmid F</sub> | 1 | 0.27 |
| IncR:Inc<sub>pA1763</sub>-KPC:IncN1 | 2 | 0.53 |
| IncP-6 | 3 | 0.80 |
| IncC | 2 | 0.53 |
| IncC:IncR | 1 | 0.27 |
| IncN1 | 1 | 0.27 |
| IncFIA<sub>pBK30661</sub> | 2 | 0.53 |
| Inc<sub>pA1763</sub>-KPC | 2 | 0.53 |
| Inc<sub>pHS062105-3</sub> | 2 | 0.53 |
| IncFIB<sub>pSC138</sub> | 1 | 0.27 |

\* These 377 plasmids came from a total of 375 cpKP isolates. There were two different *bla*<sub>KPC</sub>-carrying plasmids in each of the two following isolates: an IncFII<sub>pHN7A8</sub>:IncR plasmid plus an IncX6 one in the G030 isolate, and an IncFII<sub>R100</sub>:IncFIA<sub>pBK30661</sub> plasmid plus an IncR:IncFIB<sub>plasmid F</sub> one in the G300 isolate.

**Table S5.** The Inc group of blaKPC-carrying plasmids of the 38 complete ST11 KP genome downloaded from NCBI.

| **Inc group** | **Number** | **Percent (%)** |
| --- | --- | --- |
| IncFII<sub>pHN7A8</sub> | 3 | 7.5 |
| IncFII<sub>pHN7A8</sub>:IncN1 | 1 | 2.5 |
| IncFII<sub>pHN7A8</sub>:IncR | 27 | 67.5 |
| IncFII<sub>pHN7A8</sub>:IncR:IncN1 | 1 | 2.5 |
| IncFII<sub>pHN7A8</sub>:Inc<sub>pA1763</sub>-KPC | 1 | 2.5 |
| IncFII<sub>pHN7A8</sub>:IncΔR | 1 | 2.5 |
| IncFII<sub>K</sub>:IncR | 2 | 5 |
| IncFII<sub>K</sub>:IncR:Inc<sub>pA1763</sub>-KPC | 1 | 2.5 |
| IncR | 3 | 7.5 |

**Table S6.** Percentage of cpKP isolates resistant to the 21 antibiotics in China*.

| **Class** | **Antibiotics** | **Percentage (%)** |
| --- | --- | --- |
| Penicillins | Ampicillin | 99.8538 |
|  | Ampicillin/sulbactam | 0.956098 |
|  | Piperacillin | 0.990834 |
|  | Piperacillin/tazobactam | 0.925744 |
| Cephalosporins, first generation | Cefazolin | 0.991408 |
| Cephalosporins, second generation | Cefuroxime | 0.982398 |
|  | Cefotetan | 0.93741 |
| Cephalosporins, third generation | Ceftazidime | 0.962959 |
|  | Ceftriaxone | 0.945567 |
| Cephalosporins, fourth generation | Cefepime | 0.891937 |
| Monobactams | Aztreonam | 0.919133 |
| Carbapenems | Imipenem | 0.914203 |
|  | Meropenem | 0.86514 |
| Aminoglycosides | Amikacin | 0.494484 |
|  | Gentamicin | 0.647221 |
|  | Tobramycin | 0.715353 |
| Fluoroquinolones | Ciprofloxacin | 0.782454 |
|  | Levofloxacin | 0.681667 |
| Furanes | Macrodantin | 0.957 |
| Sulfanilamides | Sulfamethoxazole/trimethoprim | 0.702209 |

\*Data are derived from all the 31 literatures published since 2017(1-31).

**Table S7.**
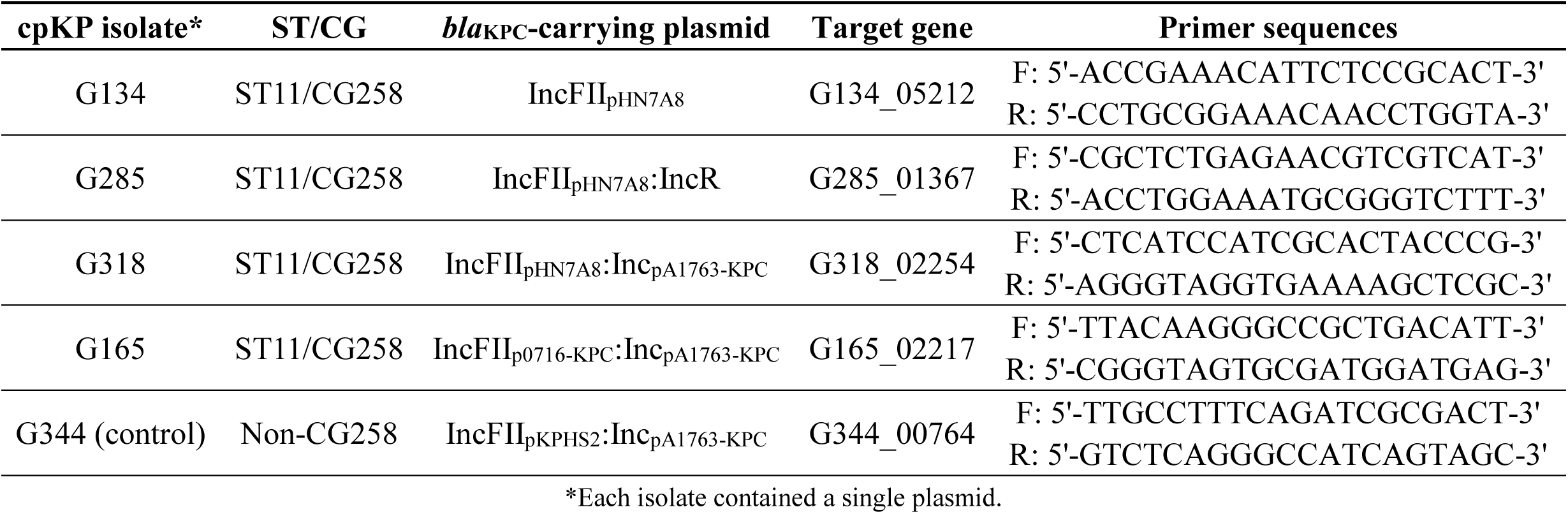
PCR primers used in the competition experiment.

**Figure S1.**
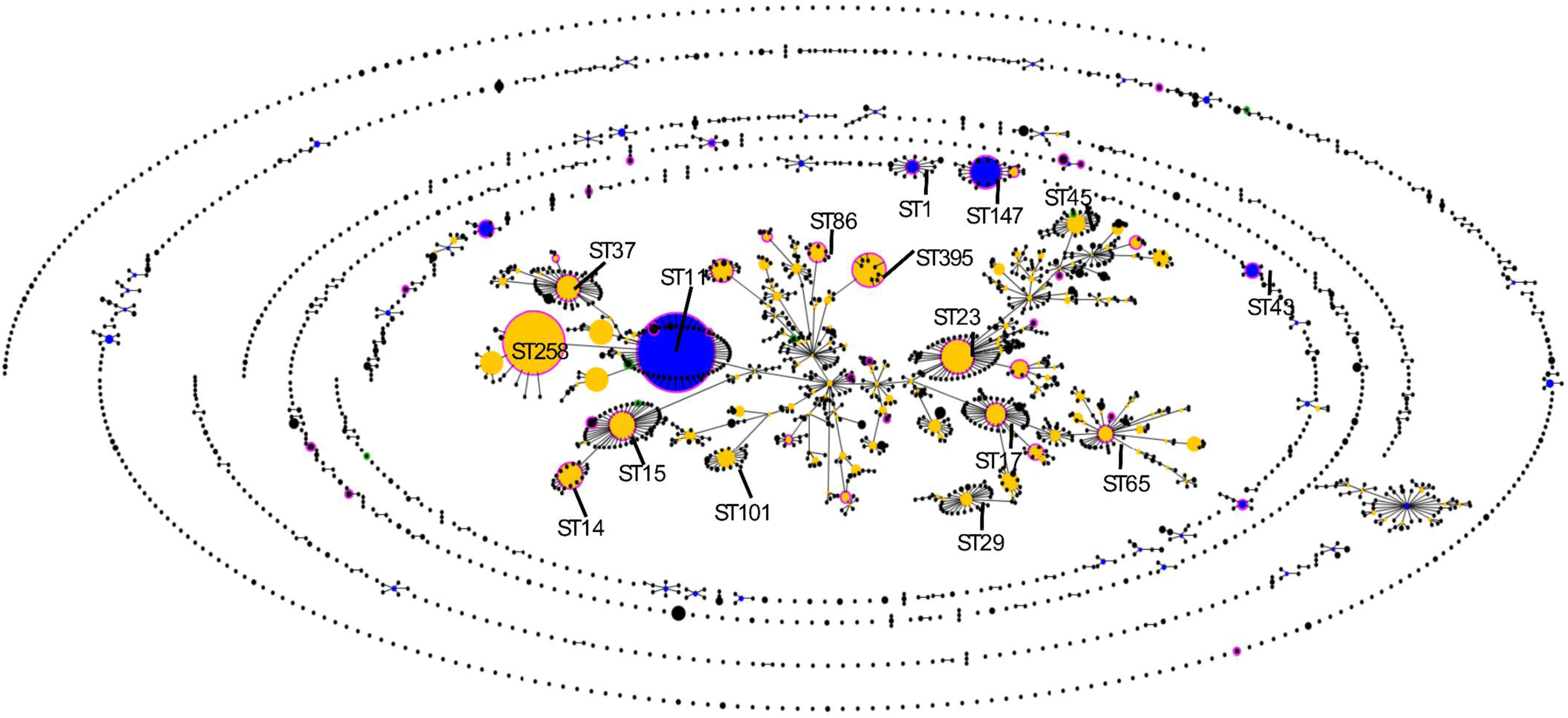
An population snapshot of the 5,099 global KP isolates. These isolates included our 420 cpKp isolates and the 4,679 isolates from the *Klebsiella* MLST database. The lines between STs indicated single-locus variants. The size of circle represented number of isolates. The circles with green rings represented the eight novel STs found in our 420 cpKp isolates, while those with cyan rings stood for the STs found in both our 420 cpKp isolates and the MLST database.

**Figure S2.**
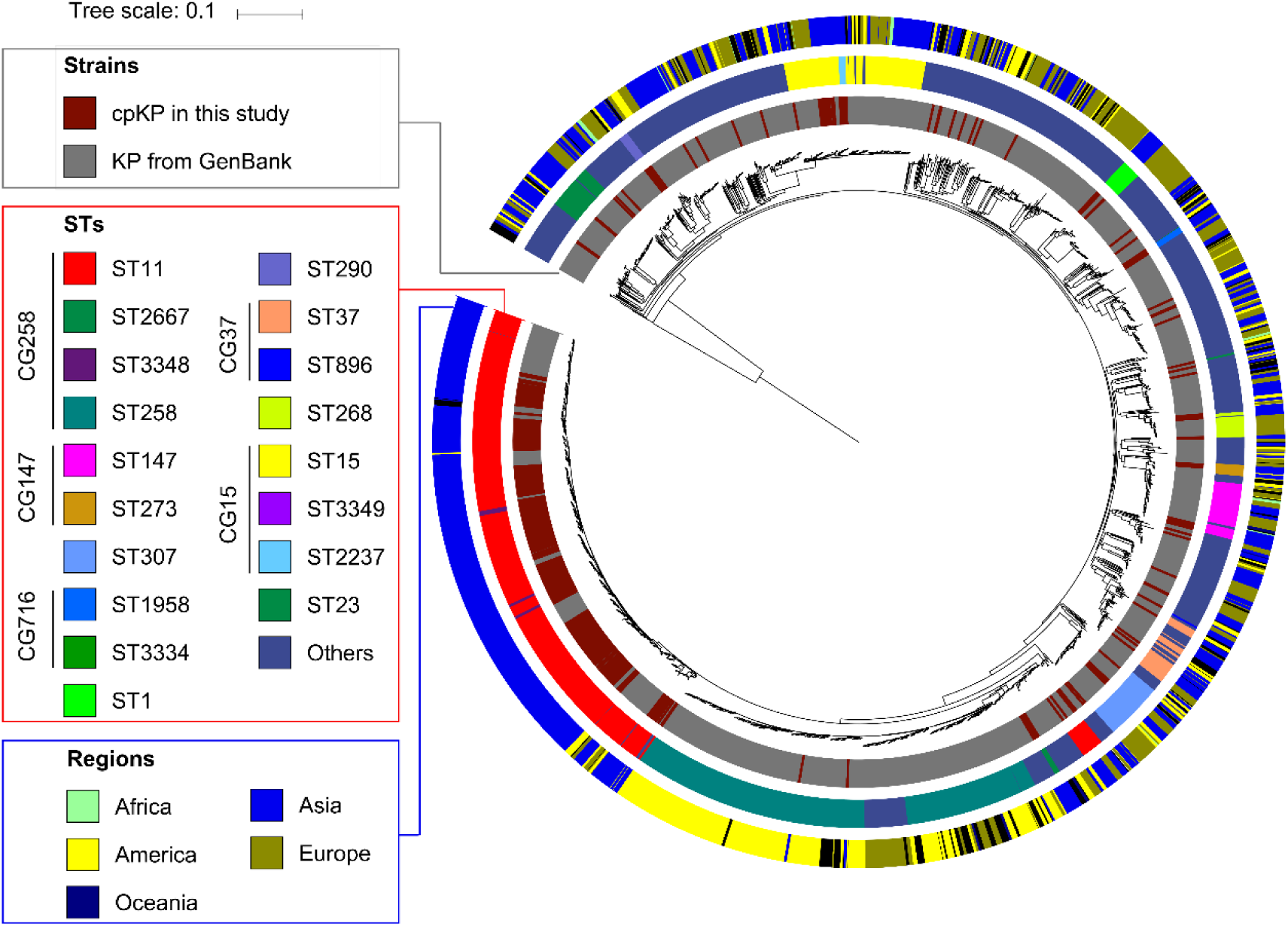
A maximum-likelihood clustering tree of the 2,300 global KP isolates. These isolates included our 420 cpKp isolates and the 1,880 isolates with determined genome sequences from GenBank (last accessed April 10, 2018). The tree was constructed from the 610,814 core SNPs of these 2,300 genome sequences. *K. variicola* DSM 15968 was used as the outgroup but not shown in the tree.

**Figure S3.**
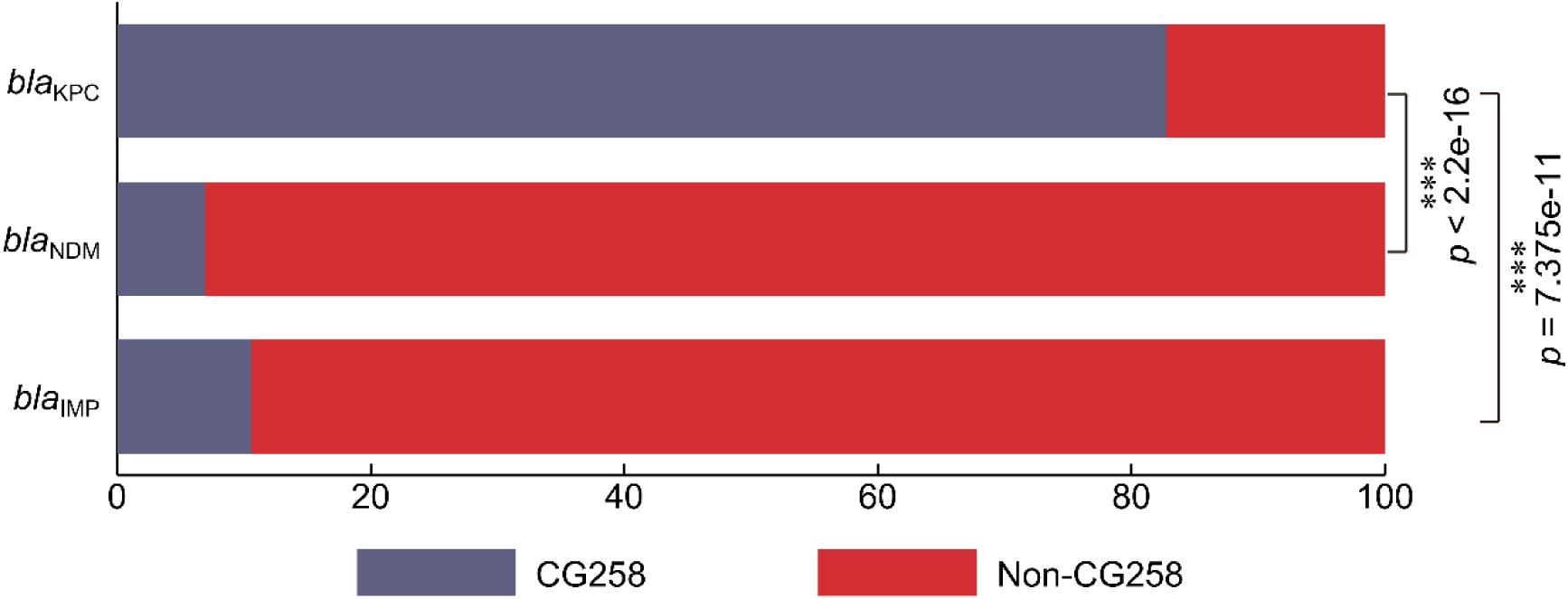
Prevalence of carbapenemase genes in CG258 and non-CG258 cpKP isolates. The numbers on the *y*-axis represented the percentage values. The *p* values were obtained using Fisher exact test. ***, statistically significant with *p* < 0.0001.

**Figure S4.**
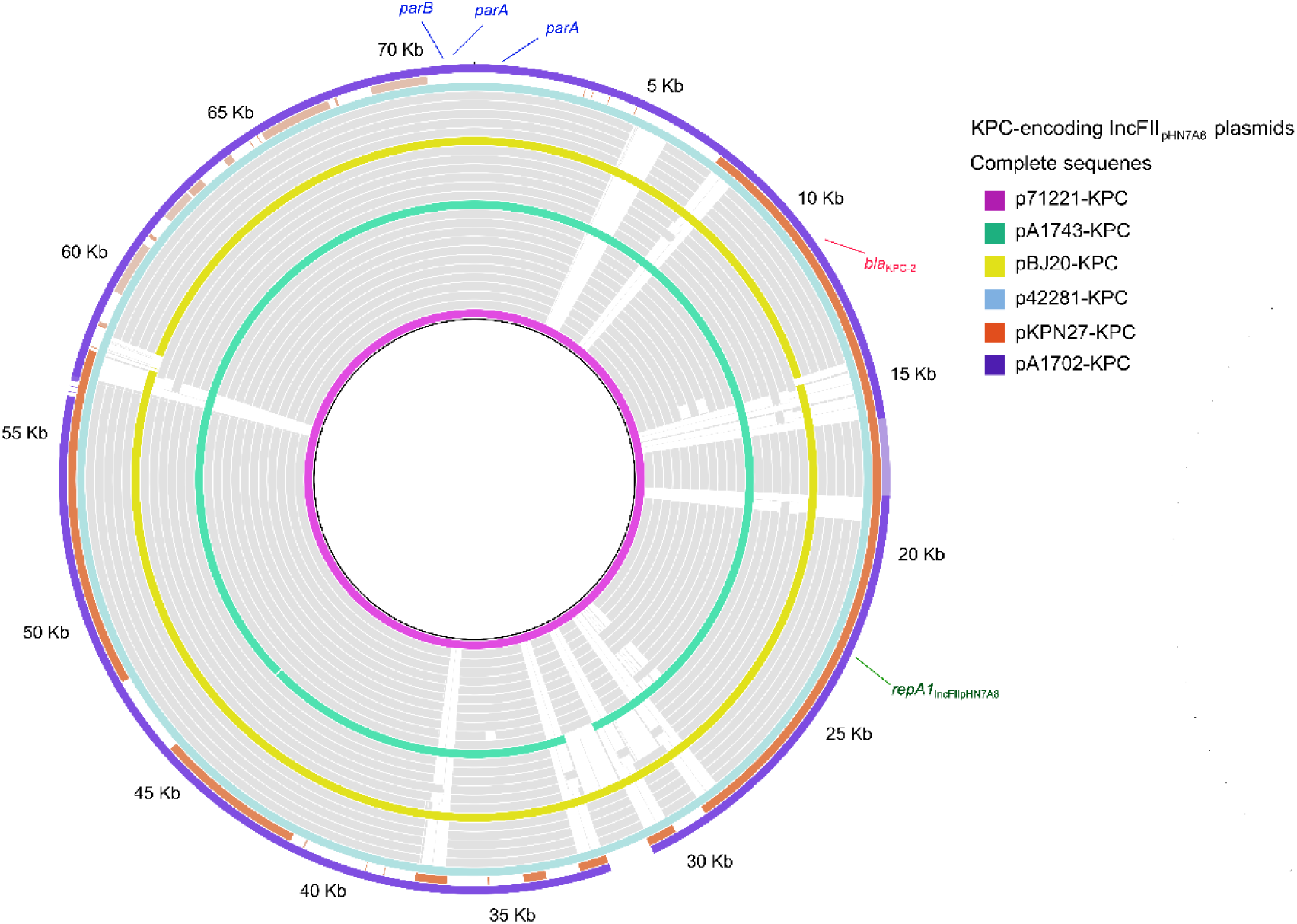
Alignment of the 28 *bla*_KPC_-carrying IncFII_pHN7A8_ plasmids from our 420 cpKp isolates. The color rings represented the fully sequenced plasmids, while the grey rings stood for those with draft sequences.

**Figure S5.**
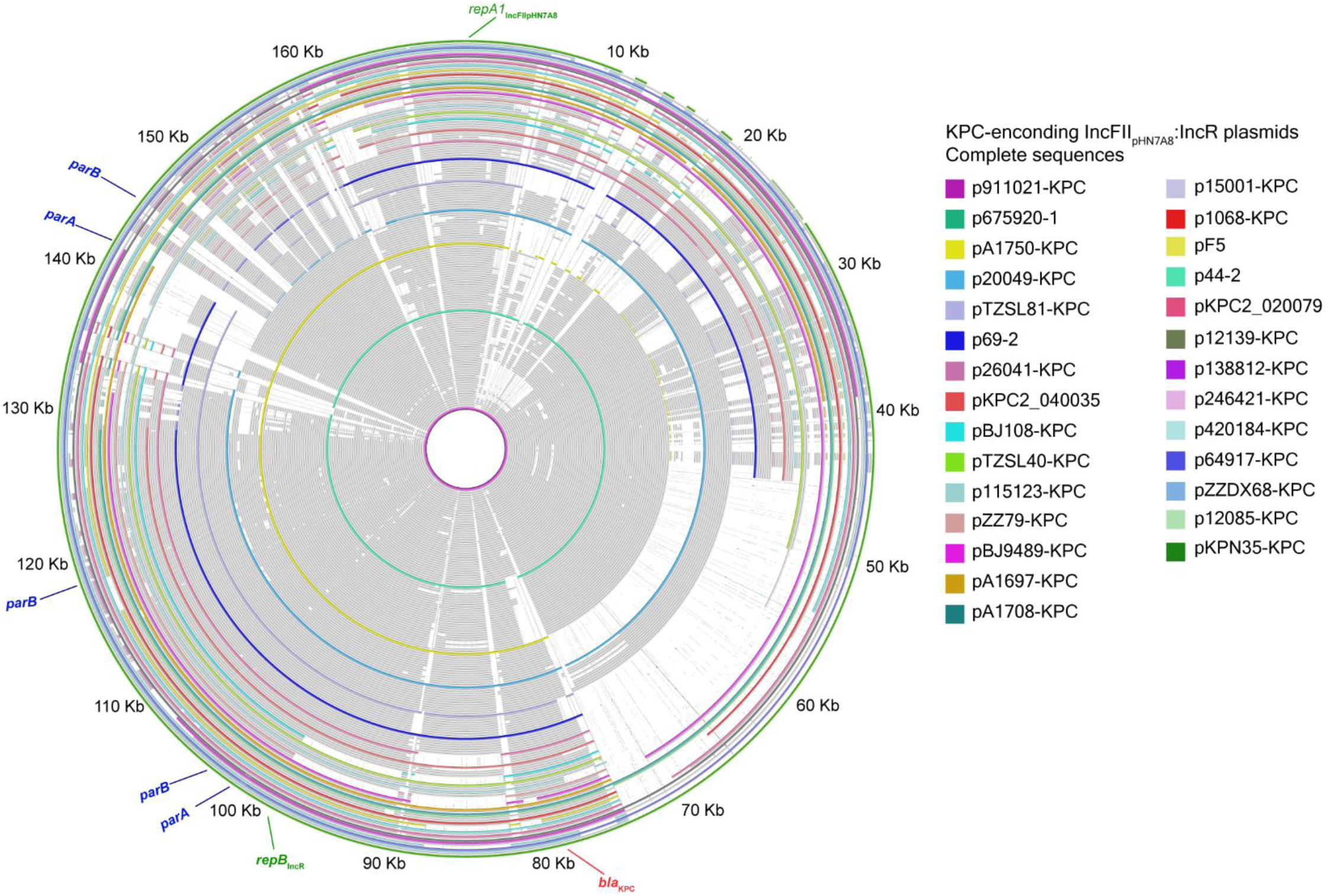
A**l**ignment **of the 163 blaKPC-carrying IncFIIpHN7A8:IncR plasmids from our 420 cpKp isolates.** The color rings represented the fully sequenced plasmids, while the grey rings stood for those with draft sequences.

**Figure S6.**
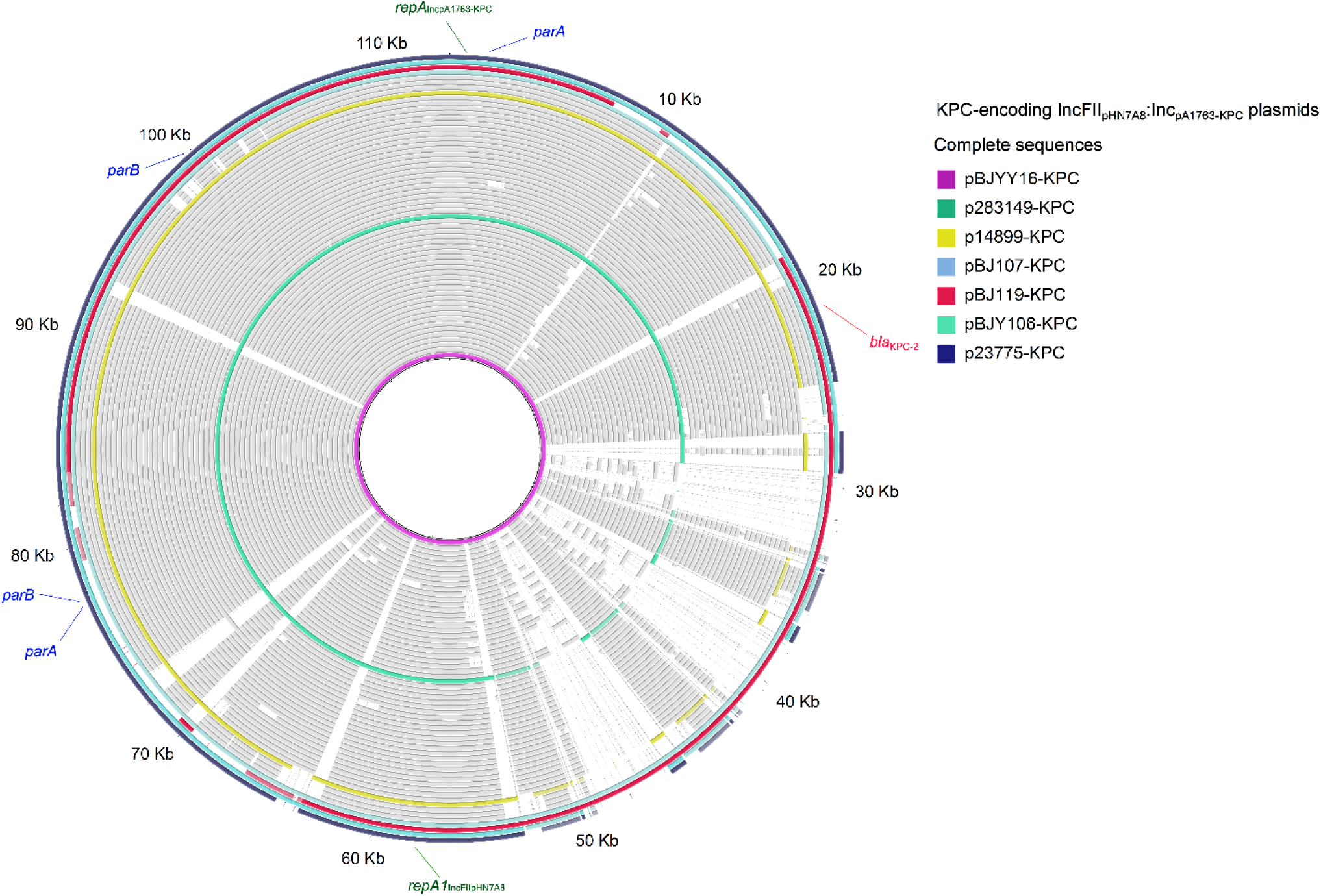
Alignment of the 59 *bla*_KPC_-carrying IncFII_pHN7A8_:Inc_pA1763-KPC_ plasmids from our 420 cpKp isolates. The color rings represented the fully sequenced plasmids, while the grey rings stood for those with draft sequences.

**Figure S7.**
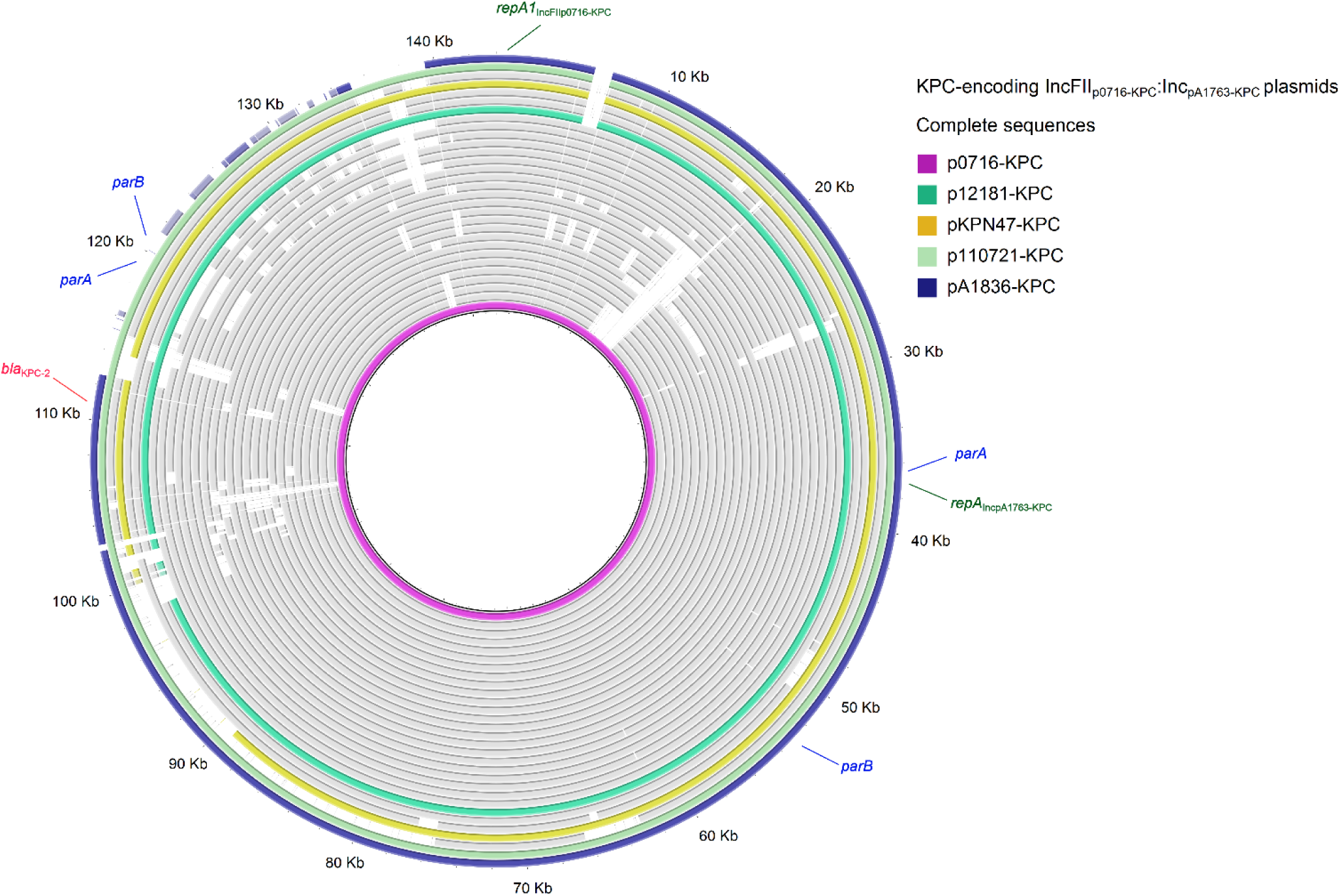
Alignment of the 30 blaKPC-carrying IncFII0716-KPC:IncpA1763-KPC plasmids from our 420 cpKp isolates. The color rings represented the fully sequenced plasmids, while the grey rings stood for those with draft sequences.

**Figure S8.**
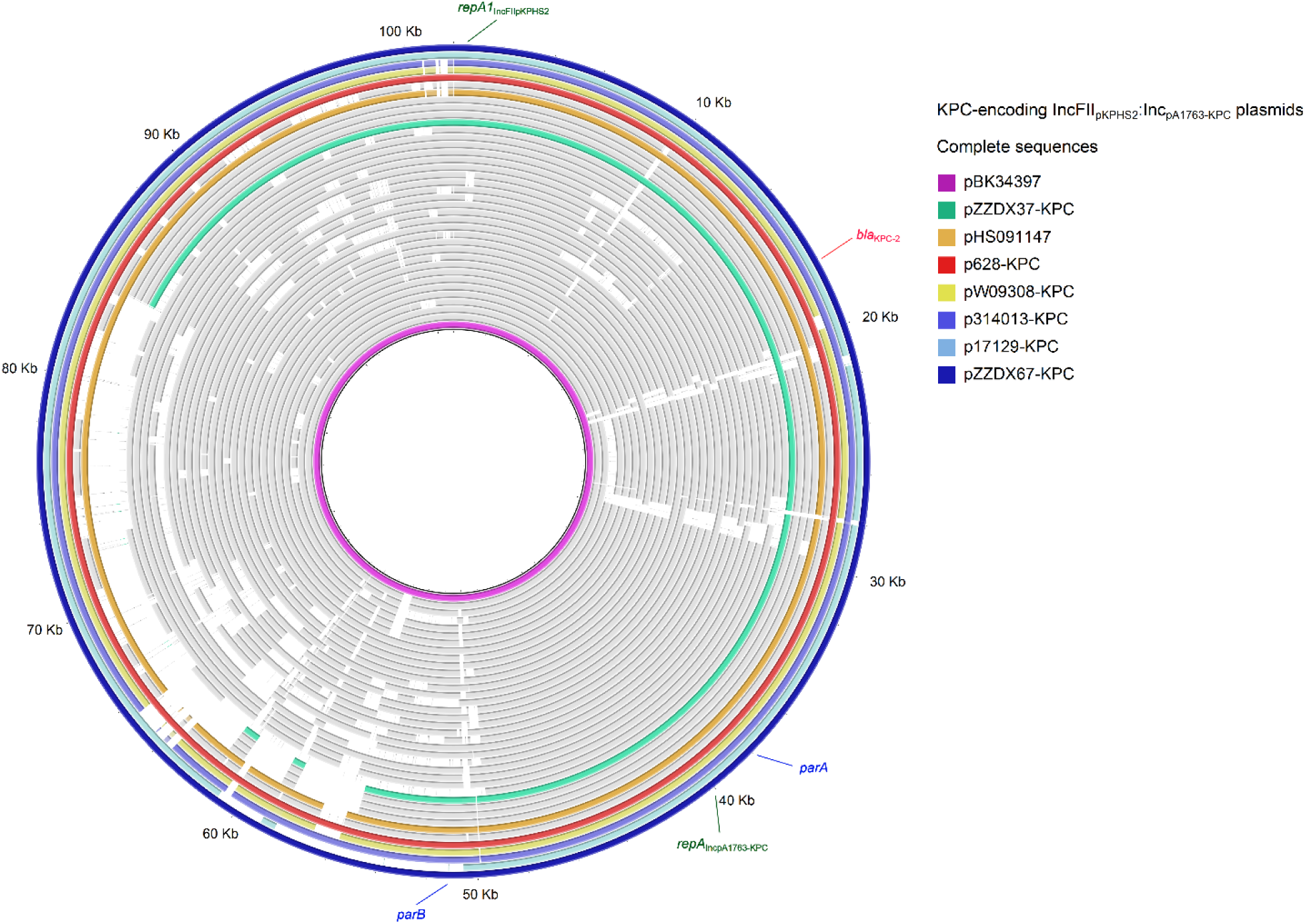
Alignment of the 36 *bla*_KPC_-carrying IncFII_pKPHS2_:Inc_pA1763-KPC_ plasmids from our 420 cpKp isolates. The color rings represented the fully sequenced plasmids, while the grey rings stood for those with draft sequences.

**Figure S9.**
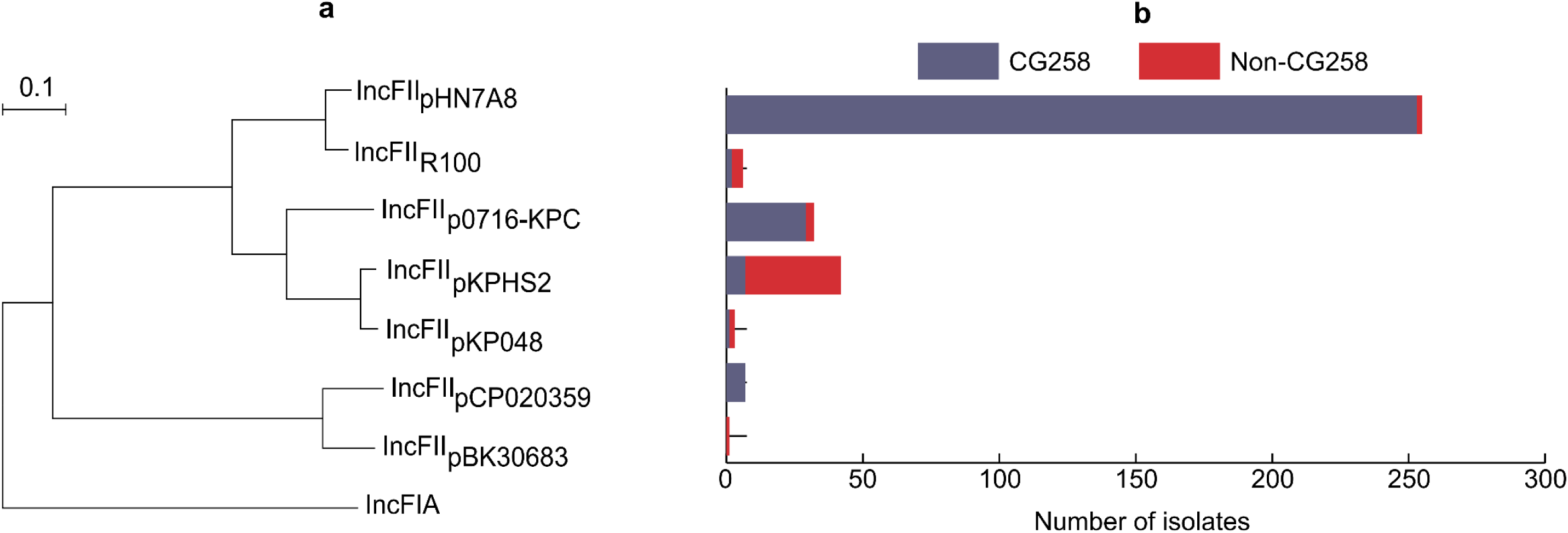
Divergence of IncFII replicons and their prevalence in CG258 and non-CG258. **a,** A maximum likelihood tree of seven subsets of IncFII replicons. The IncFIA replicon of plamid F (accession number AP001918) was used the outgroup. **b,** The prevalence of *bla*_KPC_-carrying plasmids with IncFII replicons in CG258 and non-CG258 cpKP isolates.

**Figure S10.**
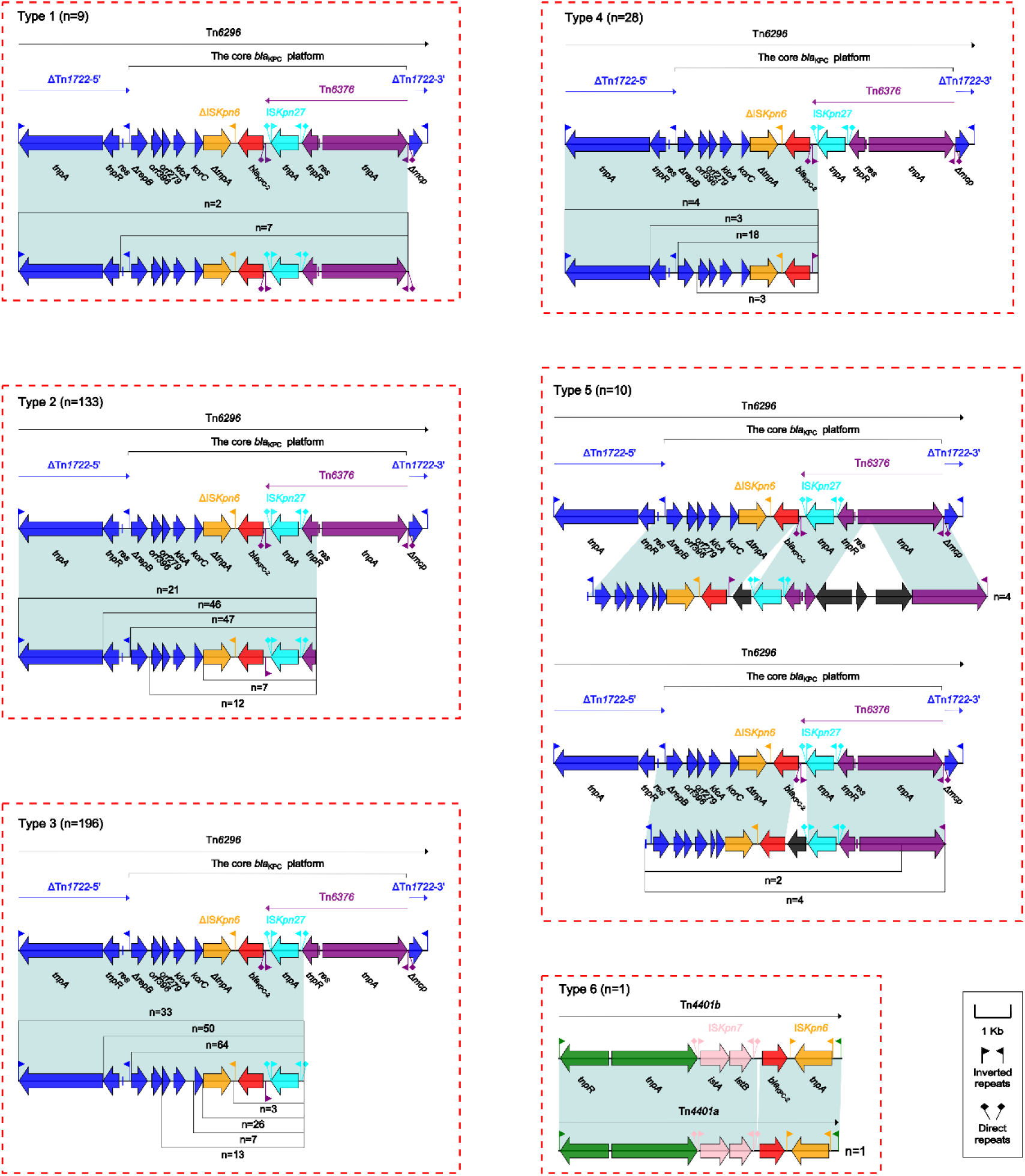
Local *bla*_KPC_ genetic environments. Genes were denoted by arrows. Genes, mobile genetic elements and other features were colored based on the functional classification. Shadow denoted homology regions (≥ 95% identity).

**Figure S11.**
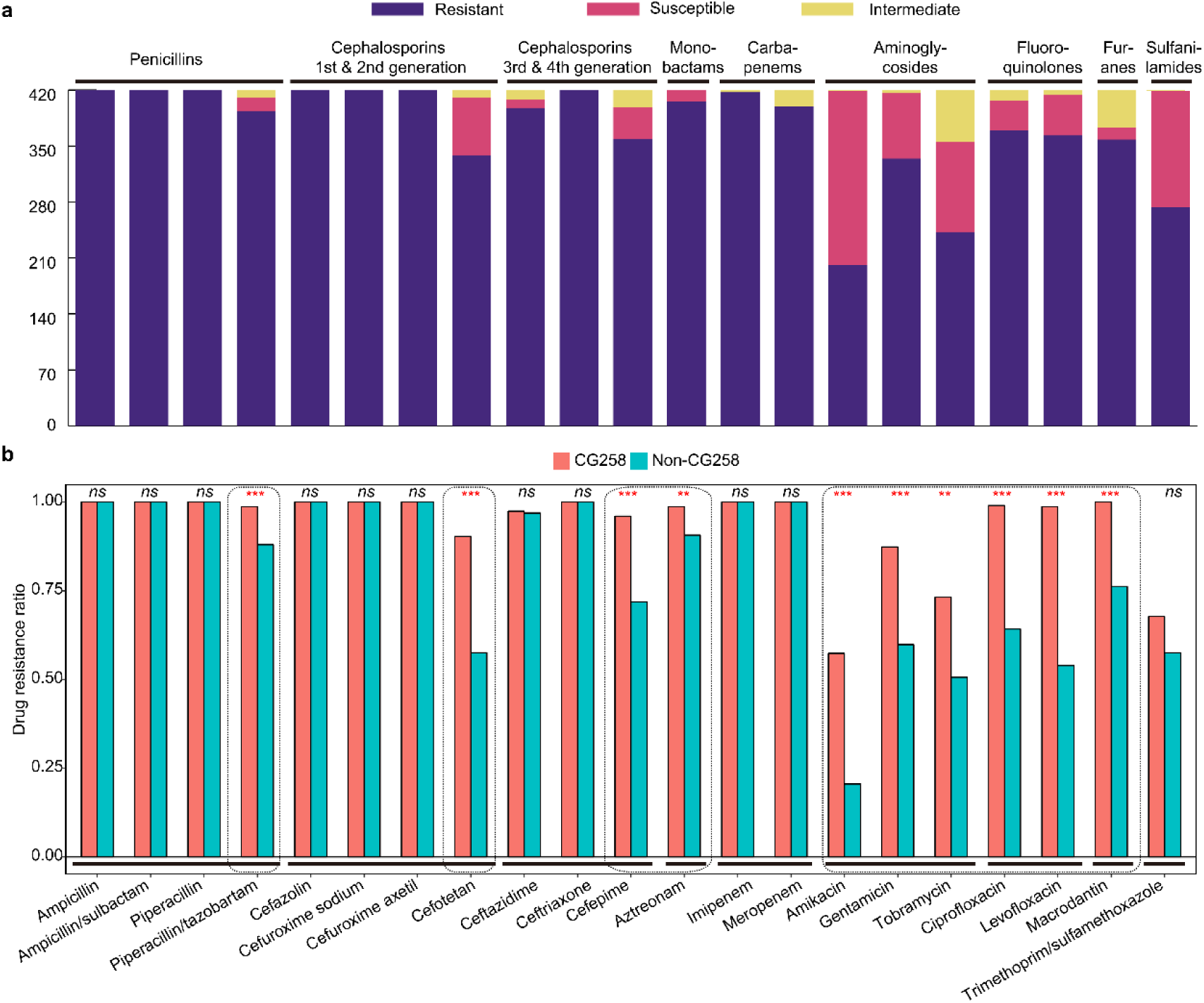
Antimicrobial susceptibility data of cpKP isolates. Shown were the drug resistance profile of our 420 cpKP isolates (a), and the resistance rates [resistant/(sensitivity + resistant)] of CG258 and non-CG258 isolates for each indicated antibiotics. A total of 21 antibiotics in nine classes were tested. The *p* values were obtained using Fisher’s exact test. NS, no significant difference. ** and ***, statistically significant with *p* < 0.001 and *p* < 0.0001, respectively.

**Figure S12.**
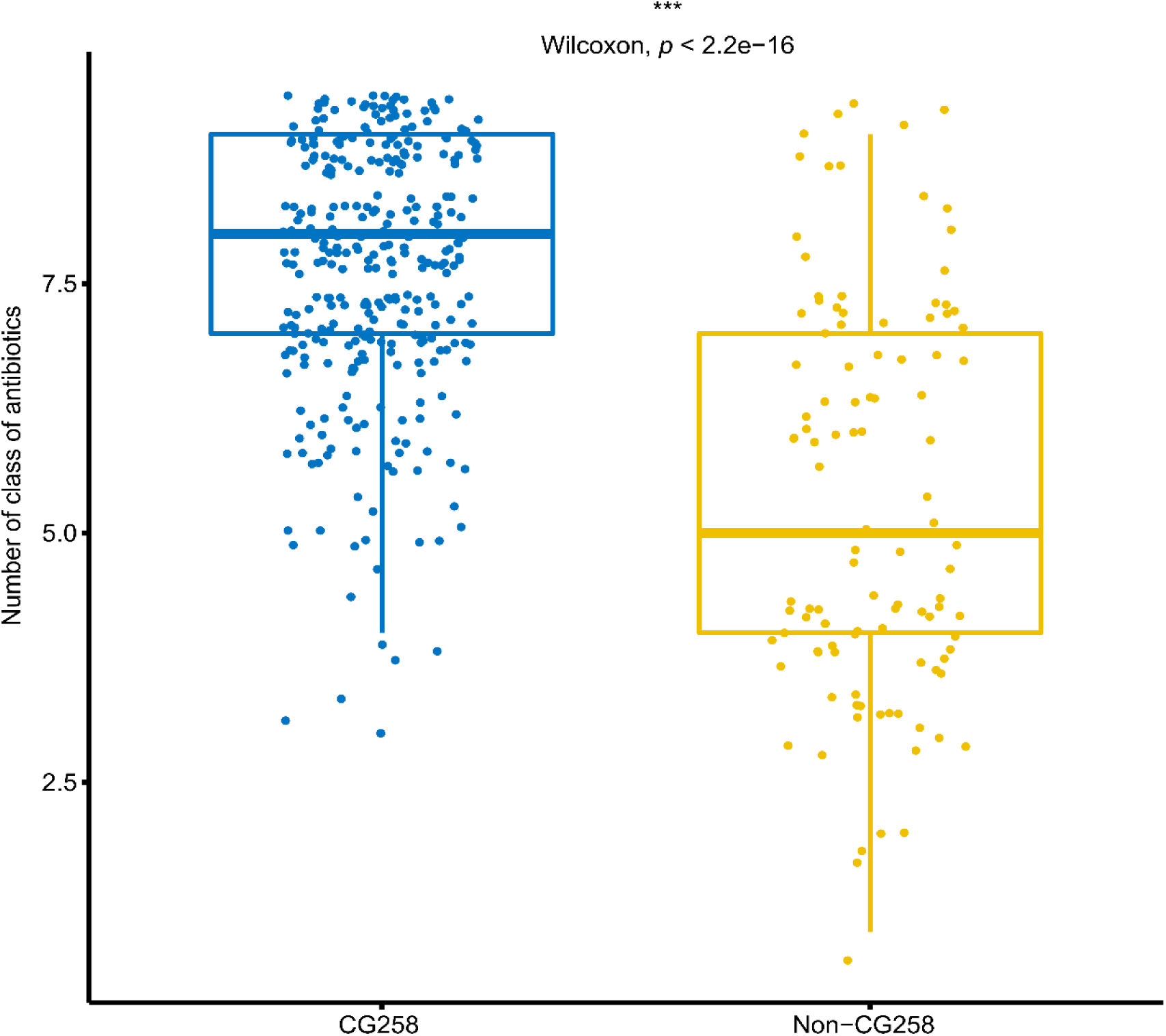
Boxplots showing the numbers of classes of antibiotics that CG258 and non-CG258 isolates were resistant to. The *p* value was tested using Wilcoxon test. ***, statistically significant with *p* < 0.0001.

**Figure S13.**
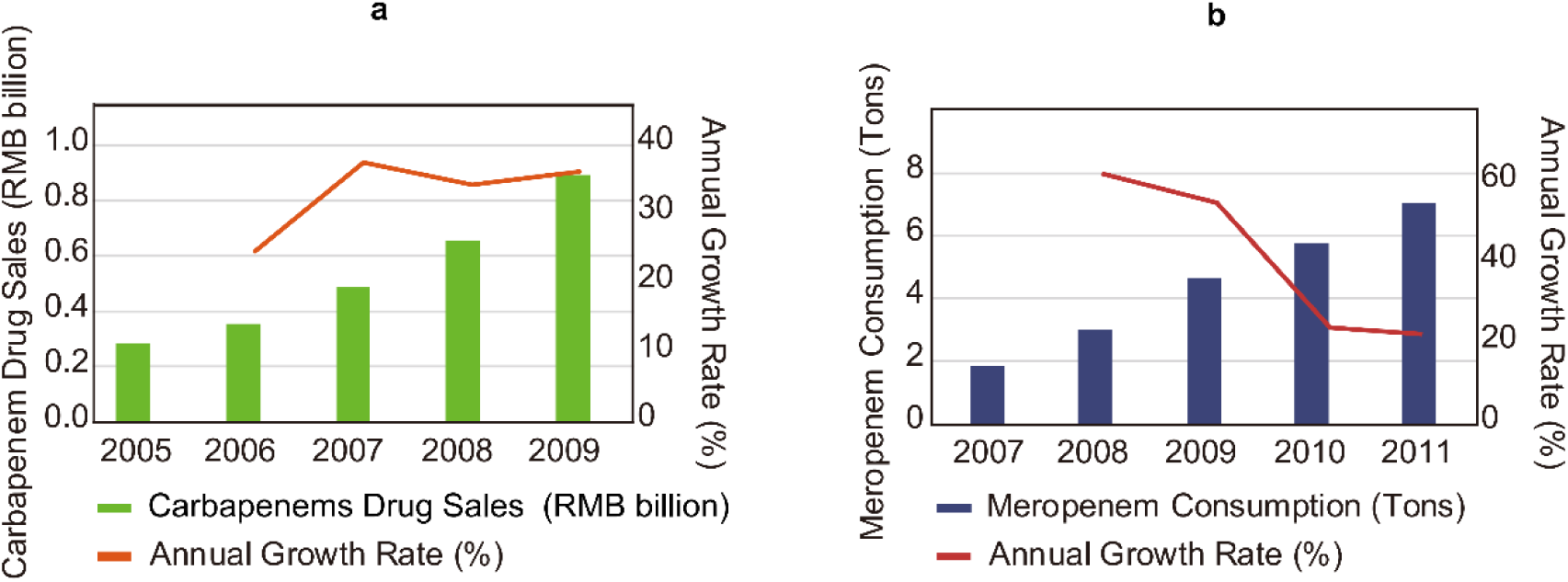
Carbapenem sales/consumption in China. **a,** Carbapenem sales in China from 2005 to 2009. **b,** Meropenem consumption in China from 2007 to 2011.

